# Mitofusin 1 and 2 regulation of mitochondrial DNA content is a critical determinant of glucose homeostasis

**DOI:** 10.1101/2021.01.10.426151

**Authors:** Vaibhav Sidarala, Jie Zhu, Elena Levi-D’Ancona, Gemma L. Pearson, Emma C. Reck, Emily M. Walker, Brett A. Kaufman, Scott A. Soleimanpour

**Affiliations:** Division of Metabolism, Endocrinology & Diabetes and Department of Internal Medicine, University of Michigan, Ann Arbor, MI 48105, USA; Vascular Medicine Institute, Division of Cardiology, Department of Medicine, University of Pittsburgh School of Medicine, Pittsburgh, PA, 15260, USA; Department of Molecular and Integrative Physiology, University of Michigan, Ann Arbor, MI 48105, USA; VA Ann Arbor Healthcare System, Ann Arbor, MI 48105, USA

## Abstract

The dynamin-like GTPases Mitofusin 1 and 2 (Mfn1 and Mfn2) are essential for mitochondrial function, which has been principally attributed to their regulation of fission/fusion dynamics. Here, we report that Mfn1 and 2 are critical for glucose-stimulated insulin secretion (GSIS) primarily through control of mitochondrial DNA (mtDNA) content. Whereas Mfn1 and Mfn2 individually were dispensable for glucose homeostasis, combined Mfn1/2 deletion in β-cells reduced mtDNA content, impaired mitochondrial morphology and networking, and decreased respiratory function, ultimately resulting in severe glucose intolerance. Importantly, gene dosage studies unexpectedly revealed that Mfn1/2 control of glucose homeostasis was dependent on maintenance of mtDNA content, rather than mitochondrial structure. Mfn1/2 maintain mtDNA content by regulating the expression of the crucial mitochondrial transcription factor Tfam, as Tfam overexpression ameliorated the reduction in mtDNA content and GSIS in Mfn1/2-deficient β-cells. Thus, the primary physiologic role of Mfn1 and 2 in β-cells is coupled to the preservation of mtDNA content rather than mitochondrial architecture, and Mfn1 and 2 may be promising targets to overcome mitochondrial dysfunction and restore glucose control in diabetes.

## INTRODUCTION

Mitochondrial dynamics, the balance of fusion and fission of mitochondrial networks, is essential to control mitochondrial structure. Mitochondrial fission and fusion are governed by several key proteins that maintain mitochondrial quality control ^1, 2^. Fission is primarily regulated by the GTPase dynamin-related protein 1 (Drp1), while fusion is controlled by both outer and inner mitochondrial membrane machinery. The dynamin-related large GTPase optic atrophy protein 1 (Opa1) controls inner membrane fusion, while outer membrane fusion is controlled by two GTPases known as Mitofusin 1 and 2 (Mfn1 and Mfn2). Mitofusin 1 and 2 share ∼80% sequence similarity and contain homologous functional domains, suggesting they play both overlapping and unique roles in metabolic function largely ascribed to their control of mitochondrial structure ^3^.

Imbalances in mitochondrial dynamics in metabolically active tissues have been implicated in many human diseases, including neurodegenerative conditions, cancer, cardiovascular disease, and diabetes ^2^. Mitochondria provide the energy necessary for β-cell insulin release ^4, 5^, and abnormalities in mitochondrial structure and bioenergetics have been observed in the β-cells of humans with type 2 diabetes (T2D; ^6, 7^). Further, mitochondrial defects have been recently found to precede the development of T2D in human β-cells ^8^. Thus, strategies to understand and overcome mitochondrial dysfunction in β-cells are of great appeal for the treatment of diabetes. However, the role of the core fission/fusion machinery in β-cells is less clear. Reduced β-cell *Mfn2* expression has been observed secondary to the deposition of toxic islet amyloid polypeptide oligomers observed in T2D as well as in mouse models of T2D ^9, 10^, yet the functions of Mfn1 and Mfn2 in β-cells *in vivo* are unexplored.

Here, we elucidate a key physiologic role for Mfn1 and 2 in the maintenance of glucose homeostasis that unexpectedly occurs through the preservation of β-cell mtDNA content, rather than mitochondrial structure. Utilizing several genetic mouse models, high resolution imaging, and transcriptomic profiling, we show that Mfn1 and 2 act in tandem to preserve β-cell mitochondrial health and, consequently, glucose-stimulated insulin secretion (GSIS). Mfn1 and 2 maintain mtDNA content by preventing loss of the mitochondrial transcription factor Tfam, which is a master regulator of mtDNA copy number control and mtRNA expression ^11, 12^. Indeed, adenoviral overexpression of Tfam ameliorates reductions in mtDNA content and GSIS in Mfn1/2-deficient β-cells, illustrating the critical importance of mtDNA copy number control by Mfn1/2 to promote glucose homeostasis.

## RESULTS

### Combined deficiency of Mfn1 and Mfn2 in β-cells leads to severe glucose intolerance and reduced glucose-stimulated insulin secretion

Mitochondrial health is vital to support β-cell insulin secretory responses to glucose or other nutrient stimuli. We hypothesized that Mfn1 and Mfn2 are required for GSIS in β-cells through their regulation of mitochondrial function. To test this hypothesis, we generated mice bearing β-cell specific deletion of Mfn1 alone (*Mfn1*^loxP/loxP^;*Ins1*-Cre), Mfn2 alone (*Mfn2*^loxP/loxP^;*Ins1*-Cre), and combined deletion of both Mfn1 and 2 (*Mfn1*^loxP/loxP^*Mfn2*^loxP/loxP^;*Ins1*-Cre), hereafter known as β-Mfn1^KO^, β-Mfn2^KO^, and β-Mfn1/2^DKO^ mice, respectively. Conditional alleles targeted the canonical G-1 GTPase motif of Mfn1 (exon 4) or Mfn2 (exon 6) (Figure S1a; ^13^). *Ins1*-Cre and floxed alone controls were phenotypically indistinguishable from one another, exhibiting no differences in body weight or glycemic control (Figure S1b-c). Therefore, they were pooled as controls for subsequent analyses. β-Mfn1^KO^, β-Mfn2^KO^, and β-Mfn1/2^DKO^ mice each exhibited an efficient reduction in Mfn1 and/or Mfn2 islet protein expression when compared to littermate controls (Ctrl; Figures 1a-b). We did not observe significant compensatory changes in Mfn2 expression in β-Mfn1^KO^ islets, nor in Mfn1 expression in β-Mfn2^KO^ islets (Figures 1a-b). Further, we did not observe differences in expression of the other core fission/fusion proteins, Drp1 and Opa1, following deletion of Mfn1 and/or Mfn2 (Figure S1d).

**Figure 1.**
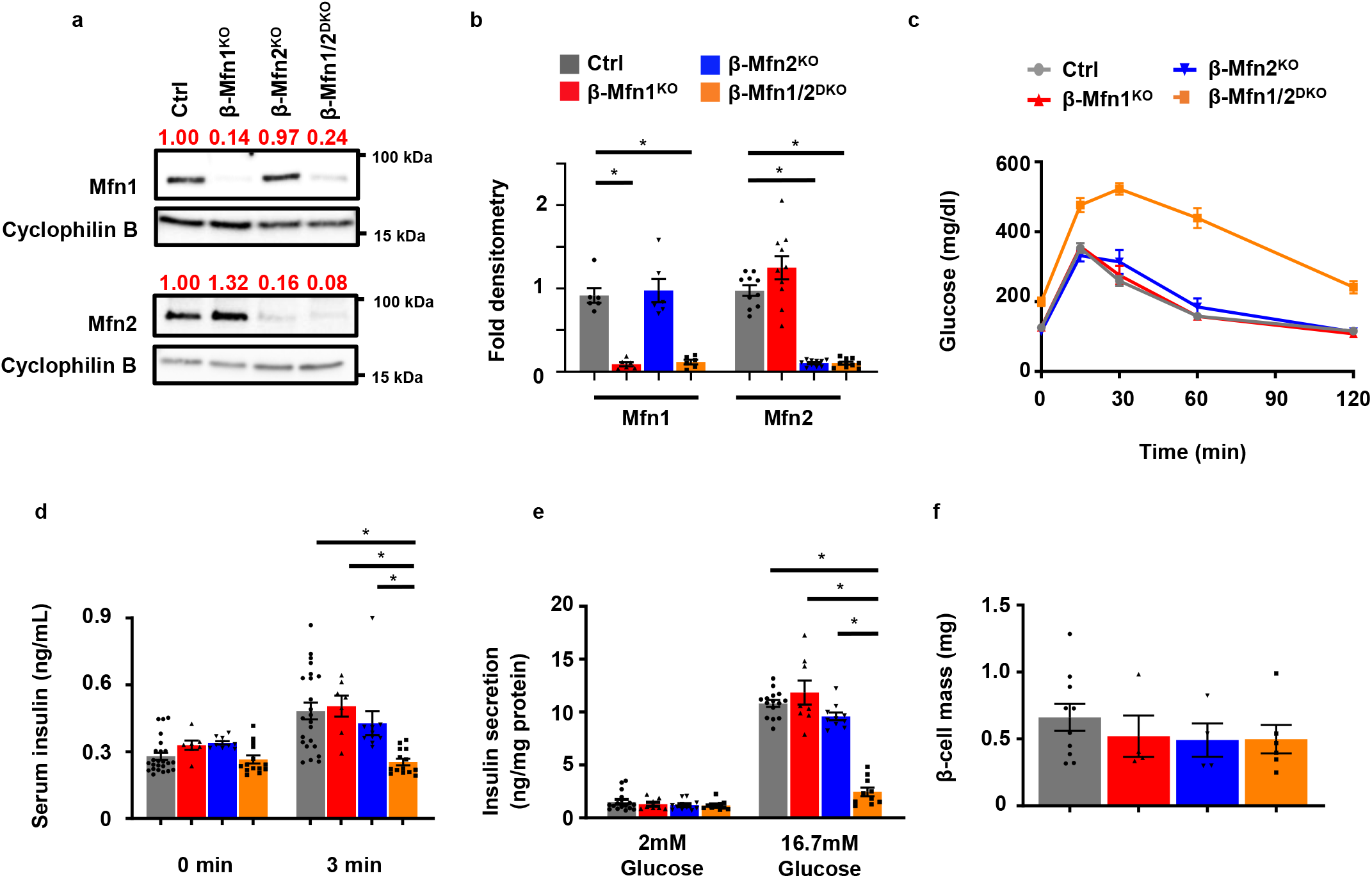
Loss of both Mfn1 and Mfn2 in β-cells impairs glucose tolerance and glucose-stimulated insulin release. **a** Expression of Mfn1 and Mfn2 by Western blot (WB) in islets isolated from 8-10-week-old Ctrl, β-Mfn1^KO^, β-Mfn2^KO^ and β-Mfn1/2^DKO^ mice. Cyclophilin B serves as a loading control. Representative of 6 independent mice/group for Mfn1 and 10 independent mice/group for Mfn2. Band intensity of each lane of displayed representative blot (shown as fold change compared to Ctrl, expression normalized to cyclophilin B) above each blot in red. **b** Densitometry (normalized to cyclophilin B) from studies in Figure 1A. n=6/group for Mfn1 and 10/group for Mfn2. Data are presented as Mean ± SEM. *p<0.0005 by one-way ANOVA followed by Tukey’s post-hoc test for multiple comparisons. **c** Blood glucose concentrations measured during IPGTT of 8-week-old Ctrl (n=28), β-Mfn1^KO^ (n=7), β-Mfn2^KO^ (n=6) and β-Mfn1/2^DKO^ (n=15) littermates. Data are presented as Mean ± SEM. (p<0.0005 by two-way ANOVA for β-Mfn1/2^DKO^ mice vs Ctrl followed by Sidak’s post-hoc test for multiple comparisons). **d** Serum insulin concentrations (n=7-24/group) measured during *in vivo* glucose-stimulated insulin release testing in 8-week-old Ctrl (n=24), β-Mfn1^KO^ (n=7), β-Mfn2^KO^ (n=10) and β-Mfn1/2^DKO^ (n=14) littermates. Data are presented as mean ± SEM. *p<0.0005 for β-Mfn1/2^DKO^ mice vs Ctrl and β-Mfn1^KO^, p=0.004 for β-Mfn1/2^KO^ vs β-Mfn2^KO^ by one-way ANOVA followed by Sidak’s post-hoc test for multiple comparisons. **e** Glucose-stimulated insulin secretion following static incubations in 2 mM and 16.7 mM glucose, performed in isolated islets of 8-week-old Ctrl (n=16), β-Mfn1^KO^ (n=8), β-Mfn2^KO^ (n=11) and β-Mfn1/2^DKO^ (n=10) littermates. Data are presented as mean from independent mice ± SEM. *p<0.0005 by one-way ANOVA followed by Sidak’s post-hoc test for multiple comparisons. **f** Pancreatic β-cell mass measured in 10-week-old Ctrl (n=10), β-Mfn1^KO^ (n=4), β-Mfn2^KO^ (n=4) and β-Mfn1/2^DKO^ (n=6) littermates. Data are presented as mean ± SEM. Uncropped western blots and source data are provided as a Source Data file.

While loss of either Mfn1 or Mfn2 alone has been shown to impair glucose homeostasis in other metabolic tissues ^3, 14^, we were surprised to find that both β-Mfn1^KO^ and β-Mfn2^KO^ mice exhibited normal glucose homeostasis during an intraperitoneal glucose tolerance test (IPGTT; Figure 1c). We also observed no changes in insulin secretion in single β-Mfn1^KO^ or β-Mfn2^KO^ mice after glucose administration *in vivo* (Figure 1d) or in isolated islets (Figure 1e). However, combined deletion of both Mfn1 and Mfn2 resulted in severe glucose intolerance (Figure 1c; p <0.005 by one-way ANOVA vs. Ctrl), due to markedly reduced GSIS (Figures 1d-e and S1e).

To evaluate other etiologies of glucose intolerance in β-Mfn1/2^DKO^ mice, we first measured β-cell mass. We observed no significant differences in β-cell mass between genotypes (Figure 1f). There were also no differences in body weight or peripheral insulin sensitivity between the groups (Figures S1f-g). Taken together, these studies suggest that Mfn1 and 2 are individually dispensable for glycemic control and GSIS, but together play vital complementary roles for the maintenance of glucose tolerance and β-cell insulin release.

### Mfn1 and 2 regulate β-cell mitochondrial structure and respiratory function

Given the connections between mitochondrial function and dynamics, we asked whether mitochondrial respiration and architecture would be defective in the islets of β-Mfn1/2^DKO^ mice, as has been reported in other models of Mfn1/2-deficiency ^15–19^. Indeed, islets isolated from β-Mfn1/2^DKO^ mice displayed a reduced glucose-stimulated oxygen consumption rate (OCR; Figures 2a and S2a) when compared to Ctrl islets. We did not observe differences in glycolysis as measured by extracellular acidification rate (ECAR; Figure S2a). Differences in glucose-stimulated OCR between Ctrl and β-Mfn1/2^DKO^ islets were no longer observed following exposure to the Complex III inhibitor antimycin A, suggesting that Mfn1/2-deficient islets primarily possess a defect in β-cell mitochondrial respiration (Figures 2a and S2a).

**Figure 2.**
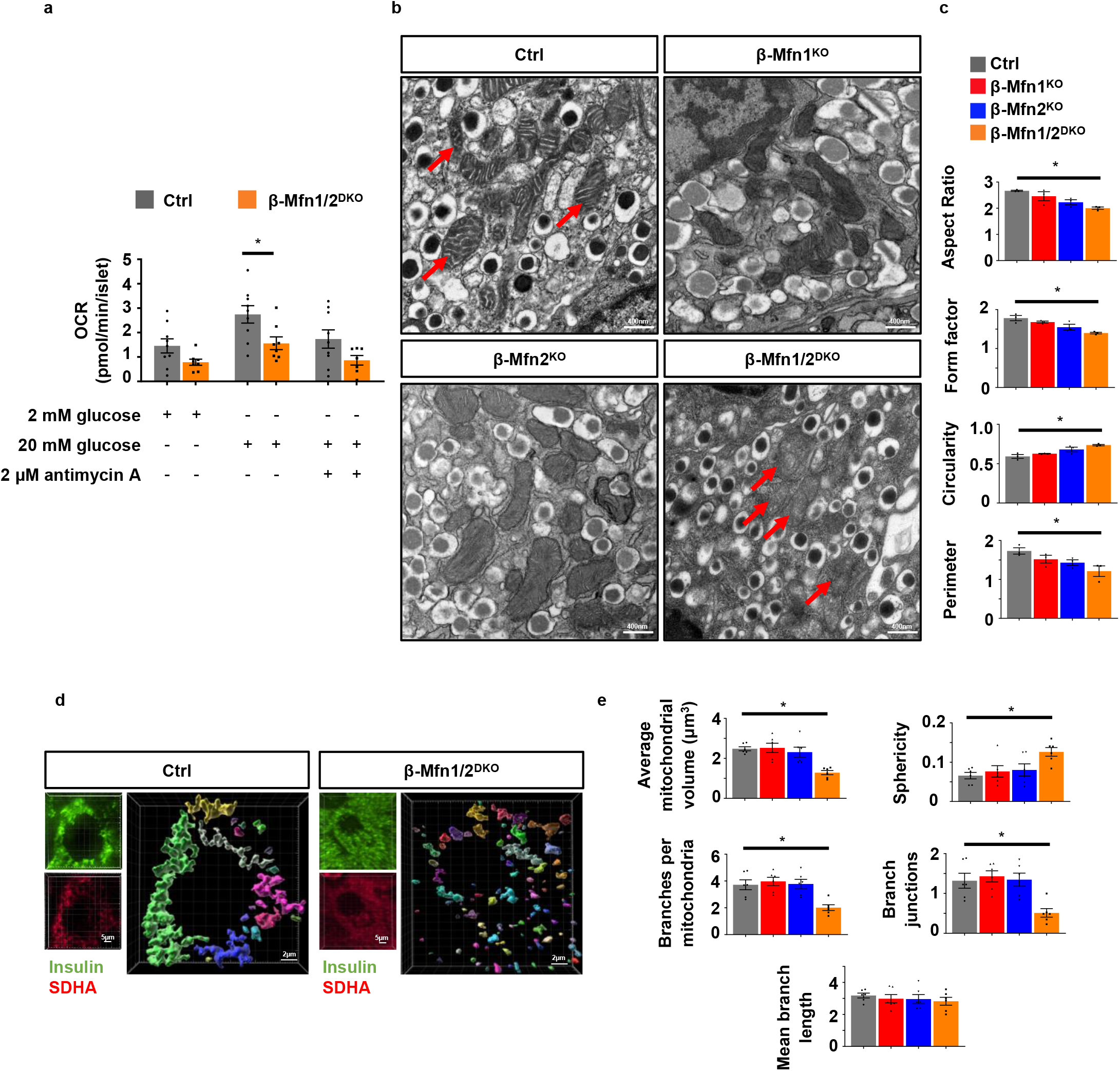
Combined Mfn1 and Mfn2 deficiency impairs mitochondrial respiration and structure in β-cells. **a** OCR measured following exposure to 2mM glucose, 20mM glucose, and 2µM antimycin A in islets isolated from 8-10-week-old Ctrl (n=9) and β-Mfn1/2^DKO^ (n=8) mice by a Seahorse flux analyzer. Data are presented as mean from independent mice ± SEM. *p=0.0176 by one-way ANOVA followed by Sidak’s post-hoc test for multiple comparisons. **b** Representative transmission EM images of β-cells from 12-week-old Ctrl, β-Mfn1^KO^, β-Mfn2^KO^ and β-Mfn1/2^DKO^ littermates. n=3/group. Mitochondria are highlighted with red arrows. **c** Quantification of mitochondrial morphology in Ctrl, β-Mfn1^KO^, β-Mfn2^KO^ and β-Mfn1/2^DKO^ β-cells from transmission EM images (∼125 independent mitochondria scored/animal). n=3/group. Data are presented as mean ± SEM. *p=0.0087 for Aspect Ratio, p=0.004 for Form factor and Circularity, p=0.037 for Perimeter by one-way ANOVA followed by Tukey’s post-hoc test for multiple comparisons. **d** Imaris® generated three-dimensional reconstruction of deconvolution immunofluorescence Z-stack images at 100X magnification stained for SDHA (see inset image – red) from pancreatic sections of Ctrl and β-Mfn1/2^DKO^ mice. β-cells were identified by insulin co-staining (inset: insulin – green). Each unique color represents a separate β-cell mitochondrial network cluster. Representative image of 6 independent mice/group. **e** β-cell mitochondrial morphology and network analysis of deconvolution immunofluorescence Z-stack images at 100X magnification stained for SDHA (and insulin) from pancreatic sections of Ctrl, β-Mfn1^KO^, β-Mfn2^KO^ and β-Mfn1/2^DKO^ mice by MitoAnalyzer. n=6/group (∼150 β-cells/animal were quantified). Data are presented as mean ± SEM. *p=0.0006 for Average mitochondrial volume, p=0.0089 for Sphericity, p=0.0032 for Branches per mitochondria, p=0.0039 for Branch junctions by one-way ANOVA followed by Sidak’s post-hoc test for multiple comparisons. Source data are provided as a Source Data file.

Loss of key mediators of mitochondrial fusion disrupt mitochondrial network balance and favor increased mitochondrial fission, often leading to the appearance of increased punctate or spherical mitochondria ^20^. We thus examined mitochondrial structure in Mfn1- and/or Mfn2-deficient β-cells by several independent and complementary methods. First, we assessed β-cell mitochondrial ultrastructure by transmission electron microscopy (TEM). While β-cells from all groups had the expected appearance of electron dense insulin granules, β-cells from β-Mfn1/2^DKO^ mice had reductions in mitochondrial aspect ratio, form factor, and perimeter, and increased mitochondrial circularity (Figures 2b-c), indicating smaller, punctate mitochondria. We observed no overt differences in ER morphology and proximity to mitochondria in β-cells from control and β-Mfn1/2^DKO^ mice (Fig. S2b), however.

Secondly, we examined mitochondrial networking utilizing high-resolution deconvolution imaging of the mitochondrial marker succinate dehydrogenase A (SDHA). We initially used Imaris® imaging software to generate three-dimensional (3D) reconstructions of β-cell mitochondria to grossly evaluate mitochondrial networking ^21, 22^. Consistent with our findings using TEM, the number of 3D mitochondrial networks appeared similar between Ctrl and single β-Mfn1^KO^ or β-Mfn2^KO^ β-cells (Figure S2c). However, Mfn1/2-double deficient mice showed a dramatic increase in the frequency of smaller β-cell mitochondrial networks (Figure 2d).

Next, we analyzed 3D deconvolution imaging of β-cell SDHA using the Mitochondria Analyzer, a validated and unbiased quantification tool used to evaluate mitochondrial morphology and network/branch complexity ^22, 23^. This confirmed that measures of mitochondrial morphology (reduced mitochondrial volume and increased sphericity) and network/branch complexity (reduced branch number and branch junctions) were significantly impaired in β-Mfn1/2^DKO^ mice, commensurate with an increase in punctate/spherical mitochondria due to decreased mitochondrial fusion (Figure 2e). Together, these data demonstrate that together Mfn1 and 2 maintain β-cell mitochondrial structure and function.

### Mfn1 and 2 maintain β-cell mtDNA content

Mitochondrial DNA copy number is vital for β-cell function and glucose homeostasis ^24, 25^. Thus, we next examined whether mtDNA content was altered by loss of Mfn1 and/or Mfn2 in β-cells. We observed a small, yet significant, reduction in mtDNA content in islets of β-Mfn2^KO^ mice that was not observed in islets of β-Mfn1^KO^ mice (Figure 3a). Interestingly, mtDNA depletion substantially worsened in the islets of β-Mfn1/2^DKO^ mice (Figure 3a), suggesting that loss of Mfn1 exacerbates the depletion of mtDNA caused by Mfn2 deficiency. We next assessed expression of the 13 mitochondrially encoded genes and observed a consistent and significant reduction in mtRNA expression in β-Mfn2^KO^ islets that was exacerbated in β-Mfn1/2^DKO^ mice (Figure 3b). Concordantly, examination of protein expression of mitochondrial OXPHOS subunits by Western blot revealed reductions of the mitochondrially encoded subunits of Complex IV, mt-Co1, and Complex III, mt-Cytb, in β-Mfn2^KO^ islets and β-Mfn1/2^DKO^ islets (Figures 3c-d). We also observed significant reductions of nuclear encoded subunits of Complex I (Ndufb8) in both β-Mfn2^KO^ and β-Mfn1/2^DKO^ islets and Complex III (Uqcrc2) in β-Mfn1/2^DKO^ islets, but no significant depletion of nuclear encoded Complex II (Sdhb) and Complex V (Atp5a) (Figures 3c-d). Depletion of mtDNA content in β-Mfn1/2^DKO^ islets did not appear to be secondary to changes in mitochondrial mass, as we did not observe reductions in Atp5a (Figures 3c-d) or in the mitochondrial outer membrane protein Tom20 (Figure 3e), a common marker of mitochondrial mass ^26^. These studies reveal that both Mfn1 and 2 (and Mfn2 alone to a lesser degree) are necessary to maintain mtDNA copy number in β-cells.

**Figure 3.**
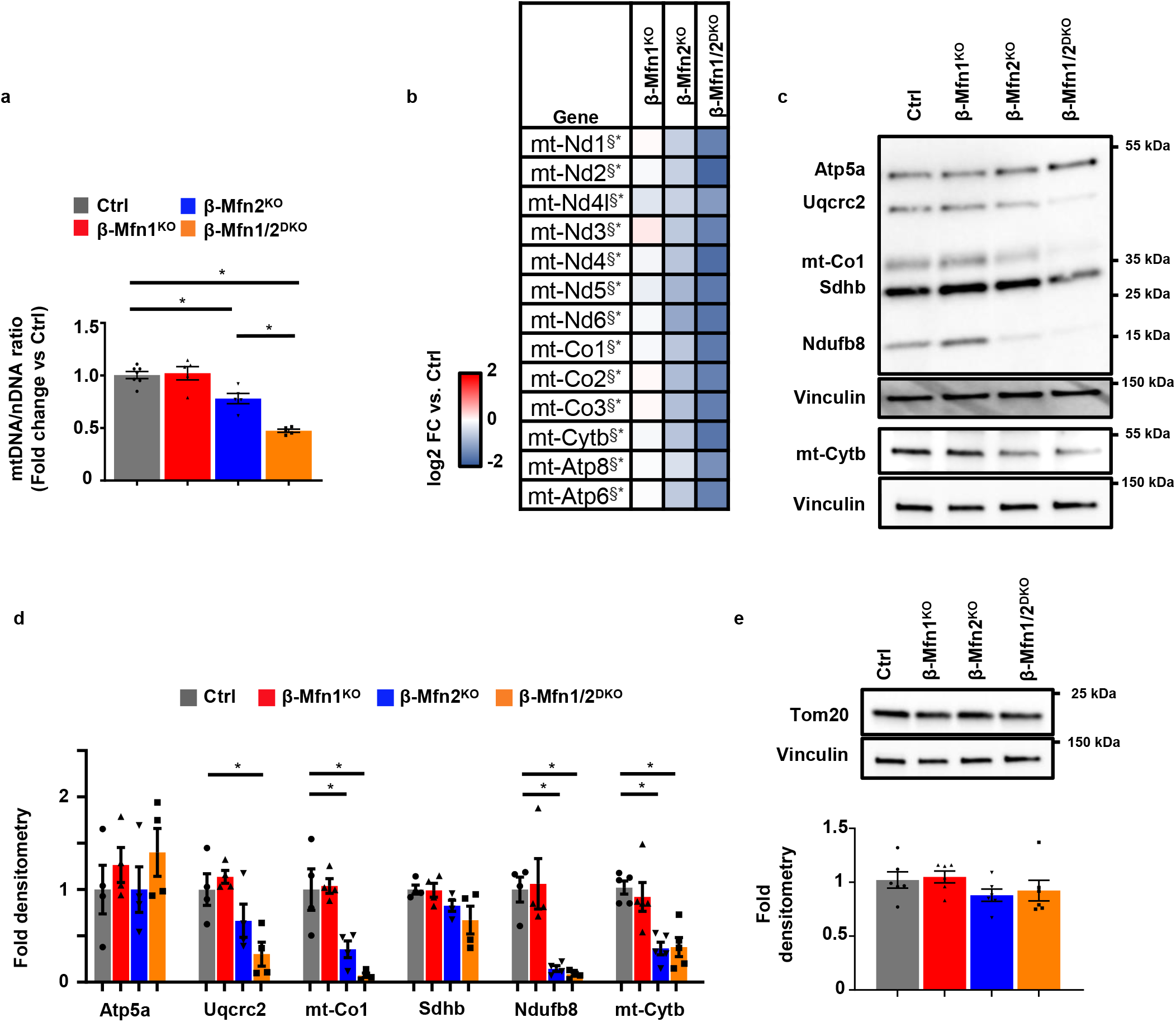
Mfn1/2-deficient β-cells exhibit reduced mtDNA content. **a** Relative mtDNA content measured by qPCR (normalized to nuclear DNA expression) in isolated islets of 8-week-old Ctrl (n=7), β-Mfn1^KO^ (n=5), β-Mfn2^KO^ (n=5) and β-Mfn1/2^DKO^ (n=5) littermates. Data are presented as mean ± SEM. *p=0.0037 for β-Mfn2^KO^ vs Ctrl, p<0.0005 for β-Mfn1/2^DKO^ vs Ctrl by one-way ANOVA followed by Sidak’s post-hoc test for multiple comparisons. **b** Heatmap representing relative expression (compared to littermate controls; Ctrl) of mitochondrial encoded transcripts in islets isolated from β-Mfn1^KO^ (n=4), β-Mfn2^KO^ (n=5) and β-Mfn1/2^DKO^ (n=5) mice. ^§*^ Benjamini-Hochberg FDR (Padj) <0.05 comparing β-Mfn2^KO^ (^§^) and β-Mfn1/2^DKO^ (*) to Ctrl. **c** Expression of OXPHOS subunit proteins by WB in islets isolated from 8-10-week-old Ctrl, β-Mfn1^KO^, β-Mfn2^KO^ and β-Mfn1/2^DKO^ mice. Representative of 4 independent mice/group. **d** OXPHOS subunit densitometry (normalized to Vinculin) from studies in Figure 3D. n=4/group; Data are presented as mean ± SEM. *p=0.02 for Uqcrc2, p<0.05 for mt-co1, p<0.001 for Ndufb8, p<0.005 for mt-Cytb by one-way ANOVA followed by Tukey’s post-hoc test for multiple comparisons. **e** Expression of Tom20 by WB in islets isolated from 8-10-week-old Ctrl, β-Mfn1^KO^, β-Mfn2^KO^ and β-Mfn1/2^DKO^ mice. Representative image (top) of 6 independent mice/group. Tom20 densitometry, normalized to vinculin (bottom). n=6/group. Data are presented as mean ± SEM. Uncropped western blots and source data are provided as a Source Data file.

### Defects in β-cell mitochondrial fusion induce glucose intolerance upon loss of mtDNA content

Our observations showing distinct and overlapping contributions of Mfn1 and 2 in the regulation of β-cell mitochondrial function led us to hypothesize that loss of mitochondrial fusion would lead to glucose intolerance due to mtDNA depletion. However, approaches to clarify the specific contribution of mtDNA depletion to metabolic dysfunction following imbalances in mitochondrial dynamics have been elusive. To this end, we generated animals harboring only a single functional allele of either *Mfn1* (β-Mfn1^+/-^Mfn2^KO^) or *Mfn2* (β-Mfn1^KO^Mfn2^+/-^) to ascertain the relative importance of mtDNA content and structure in the maintenance of glycemic control. For robustness of interpretation, these experiments were performed alongside littermate controls and β-Mfn1/2^DKO^ mice as previously presented (Figures 1-3). As expected, neither β-Mfn1^+/-^ Mfn2^KO^ nor β-Mfn1^KO^Mfn2^+/-^ mice developed changes in β-cell mass, body weight, or insulin sensitivity (Figures S3a-c). Interestingly, 3D quantification of mitochondrial morphology and network integrity revealed that, similar to β-Mfn1/2^DKO^ mice, a single allele of either *Mfn1* or *Mfn2* was not sufficient to maintain β-cell mitochondrial structure, leading to increases in punctate mitochondria consistent with loss of mitochondrial fusion (Figures 4a-b).

**Figure 4.**
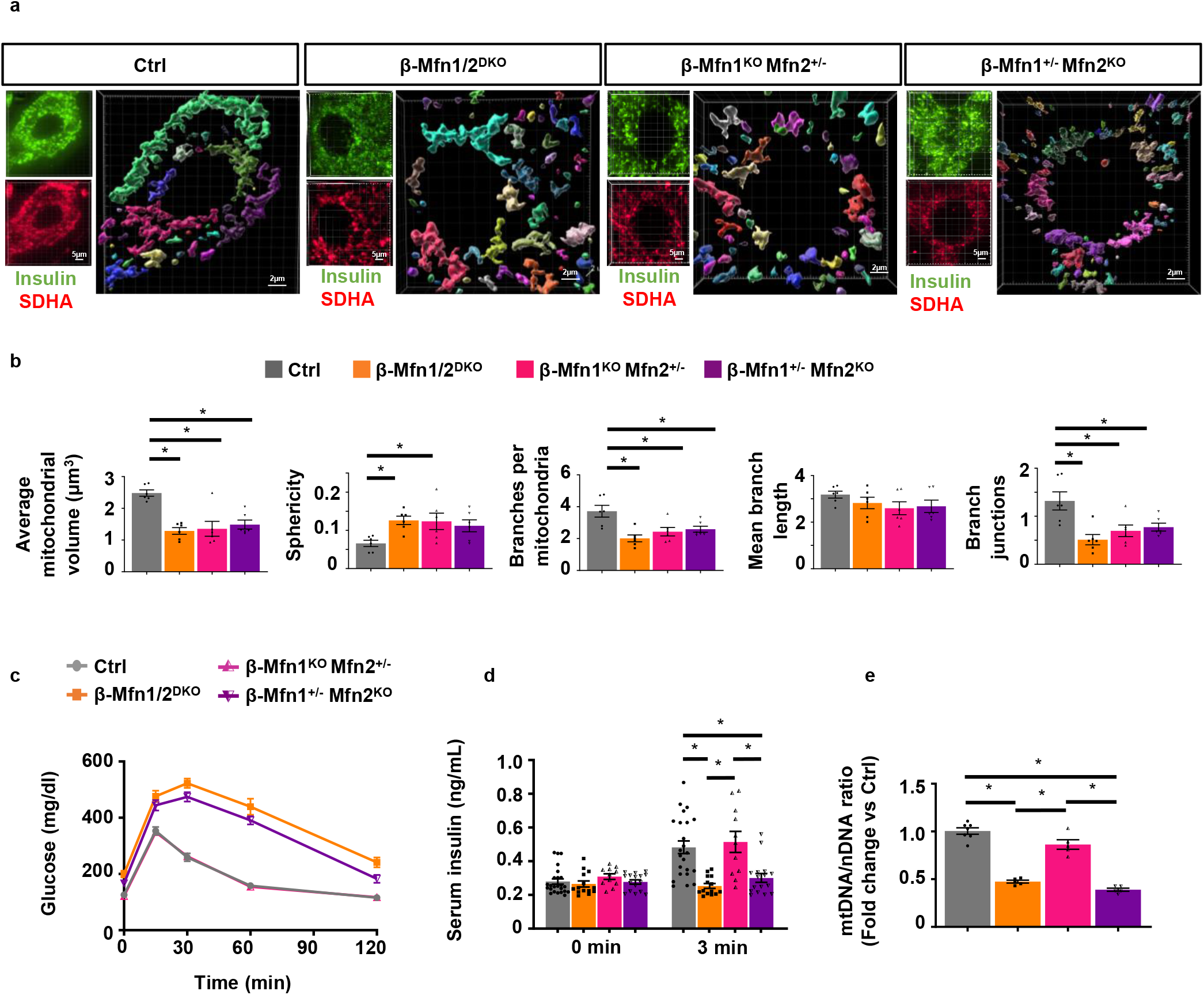
A single allele of *Mfn2*, but not *Mfn1*, maintains glycemic control by preserving β-cell mtDNA content despite impaired mitochondrial structure. **a** Imaris® generated three-dimensional reconstruction of deconvolution immunofluorescence Z-stack images at 100X magnification stained for SDHA (see inset image – red) from pancreatic sections of Ctrl, β-Mfn1/2^DKO^, β-Mfn1^KO^Mfn2^+/-^ and β-Mfn1^+/-^Mfn2^KO^ mice. β-cells were identified by insulin co-staining (inset: insulin – green). Each unique color represents a separate β-cell mitochondrial network cluster. Representative image of 6 independent mice/group. **b** β-cell mitochondrial morphology and network analysis of deconvolution immunofluorescence Z-stack images from pancreatic sections stained for insulin and SDHA of Ctrl, β-Mfn1/2^DKO^, β-Mfn1^KO^Mfn2^+/-^ and β-Mfn1^+/-^Mfn2^KO^ mice by MitoAnalyzer. n=6/group (50-200 β-cells/animal were quantified). *p=0.005 for Average mitochondrial volume, p=0.05 for Sphericity, p=0.05 for Branches per mitochondria, p=0.05 for Branch junctions by one-way ANOVA followed by Sidak’s post-hoc test for multiple comparisons. **c** Blood glucose concentrations measured during IPGTT of 8-week-old Ctrl (n=28), β-Mfn1/2^DKO^ (n=15), β-Mfn1^KO^Mfn2^+/-^ (n=13) and β-Mfn1^+/-^Mfn2^KO^ (n=17) littermates. (p<0.0005 by two-way ANOVA for β-Mfn1^+/-^Mfn2^KO^ and β-Mfn1/2^DKO^ mice vs Ctrl followed by Sidak’s post-hoc test for multiple comparisons). **d** Serum insulin concentrations (n=11-24/group) measured during *in vivo* glucose-stimulated insulin release testing in 8-week-old Ctrl (n=24), β-Mfn1/2^DKO^ (n=14), β-Mfn1^KO^Mfn2^+/-^ (n=11) and β-Mfn1^+/-^Mfn2^KO^ (n=16) littermates. *p<0.0005 by one-way ANOVA followed by Sidak’s post-hoc test for multiple comparisons. **e** Relative mtDNA content measured by qPCR (normalized to nuclear DNA expression) in isolated islets of 8-week-old Ctrl (n=7), β-Mfn1/2^DKO^ (n=5), β-Mfn1^KO^Mfn2^+/-^ (n=5) and β-Mfn1^+/-^Mfn2^KO^ (n=5) littermates. Data are presented as mean ± SEM. *p<0.0005 by one-way ANOVA followed by Sidak’s post-hoc test for multiple comparisons. Of note, studies in Ctrl and β-Mfn1/2^DKO^ mice were performed together alongside all β-Mfn1^KO^, β-Mfn2^KO^, β-Mfn1^KO^Mfn2^+/-^ (n=11) and β-Mfn1^+/-^Mfn2^KO^ (n=16) littermates and thus may appear twice for purposes of relevant comparisons (in Figures 1C, 1D, 2D, 2E, 3A and again in Figures 4A-E). Source data are provided as a Source Data file.

Despite mitochondrial morphologic imbalances in both β-Mfn1^+/-^Mfn2^KO^ and β-Mfn1^KO^Mfn2^+/-^ mice, *only* β-Mfn1^+/-^Mfn2^KO^ mice developed glucose intolerance and impaired GSIS similar to that of β-Mfn1/2^DKO^ mice (Figures 4c-d). We also found significant reductions in mtDNA content associated with the loss of GSIS observed in β-Mfn1^+/-^Mfn2^KO^ and β-Mfn1/2^DKO^ mice (Figure 4e). Thus, maintenance of a single functional allele of *Mfn2*, but not *Mfn1*, is able to preserve glucose homeostasis, mtDNA content, and insulin secretion, despite being insufficient to maintain β-cell mitochondrial network integrity. Importantly, these results disjoin β-cell function from changes in mitochondrial morphology, and implicate mtDNA copy number as the vital purveyor of glycemic control following loss of mitochondrial fusion.

### Mfn1 and 2 deficiency induces a transcriptional signature consistent with defects in mitochondrial metabolism and alterations in mtDNA replication

To begin to understand the mechanisms underlying control of mtDNA content by Mfn1 and 2, we analyzed bulk RNA sequencing (RNAseq) data generated from islets of β-Mfn1^KO^, β-Mfn2^KO^, and β-Mfn1/2^DKO^ mice as well as littermate controls. Initial hierarchical clustering analyses based on the top 500 most highly expressed genes revealed that only β-Mfn1/2^DKO^ islets clustered by genotype, while we did not observe discrete clustering across Ctrl or single β-Mfn1^KO^ and β-Mfn2^KO^ islets due to very few differences in expression between these groups (Figures 5a and S4a-b). Gene ontology and pathway analyses of differentially expressed genes from β-Mfn1/2^DKO^ islets revealed significant changes related to endocrine hormone/insulin secretion, β-cell signature genes, as well as several metabolic pathways (Figures 5b and S4c-d), consistent with the defects in β-cell function and mitochondrial metabolism observed in β-Mfn1/2^DKO^ mice.

**Figure 5.**
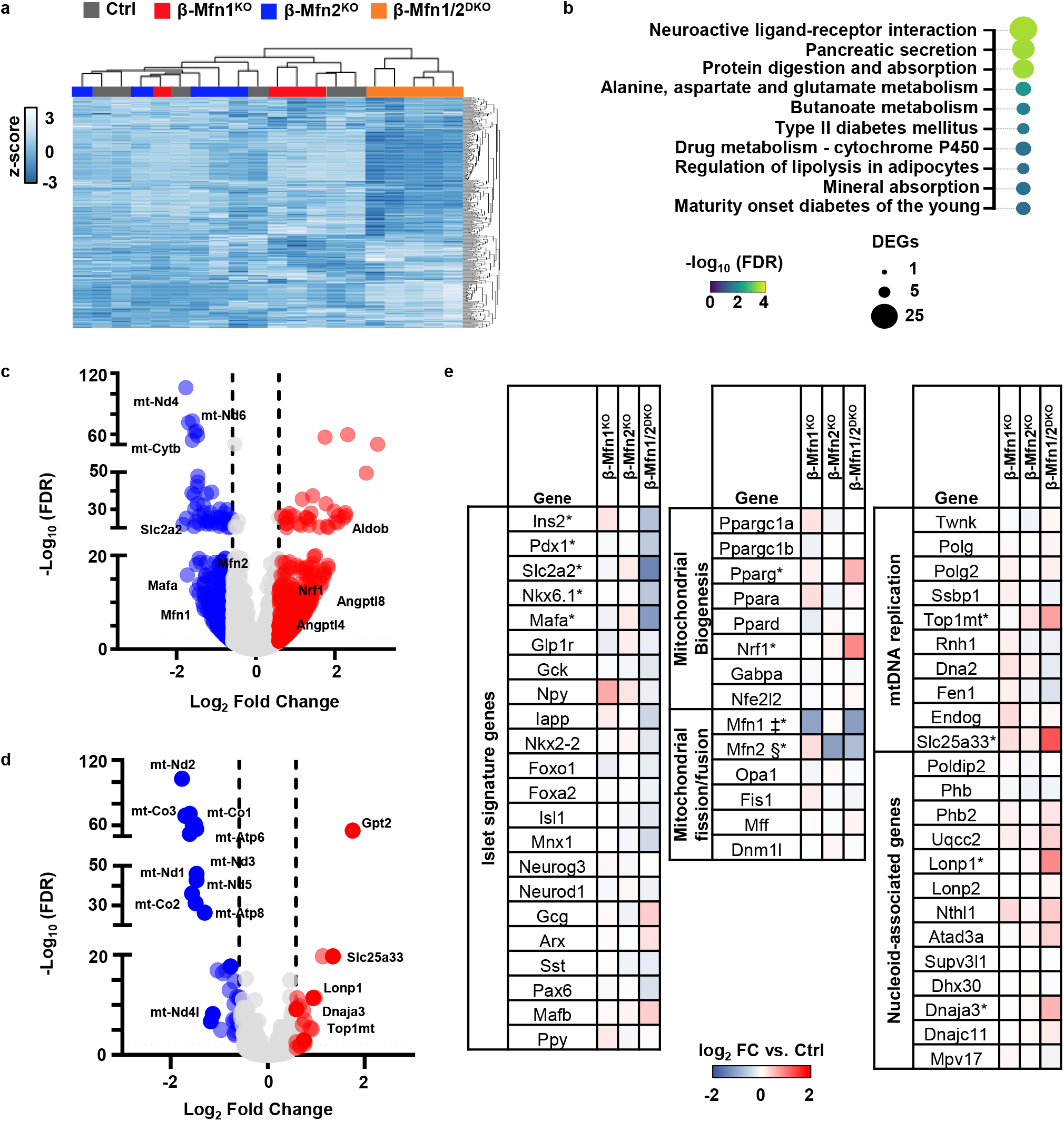
Mfn1/2-deficiency induces expression of genes associated with mtDNA replication. **a** Hierarchical clustering of RNAseq samples displaying the 500 genes with the highest mean expression in islets of 12-week-old Ctrl (n=6), β-Mfn1^KO^ (n=4), β-Mfn2^KO^ (n=5) and β-Mfn1/2^DKO^ (n=5) mice. **b** Pathway analysis of top differentially regulated pathways in β-Mfn1/2^DKO^ islets (n=5) compared to littermate Ctrl islets (n=6), demarcated by both false discovery rate (FDR) and the number of significantly differentially expressed genes (DEGs) per pathway. **c** Volcano plot depicting differential RNA expression in islets of β-Mfn1/2^DKO^ (n=5) mice compared to littermate Ctrl mice (n=6). Significantly differentially expressed genes demarcated by -log10 FDR > or < 2 and log2 fold change (FC) > or < 0.6. **d** Volcano plot depicting differential RNA expression of MitoCarta 2.0 targets in islets of β-Mfn1/2^DKO^ (n=5) mice compared to littermate Ctrl mice (n=6). Significantly differentially expressed genes demarcated by -log10 FDR > or <2 and log2 fold change (FC) > or <0.6. **e** Differential RNA expression heatmap of selected genes from islets of β-Mfn1^KO^ (n=4), β-Mfn2^KO^ (n=5) and β-Mfn1/2^DKO^ (n=5) mice compared to littermate Ctrl mice (n=6). up. ‡§* Benjamini-Hochberg FDR (Padj) <0.05 comparing β-Mfn1^KO^(‡), β-Mfn2^KO^(§) and β-Mfn1/2^DKO^(*) to Ctrl.

The nearly 1,000 significantly dysregulated genes in β-Mfn1/2^DKO^ islets included downregulation of mitochondrial encoded transcripts, as well as key β-cell signature genes such as *Slc2a2* (Glut2) and *MafA* (Figures 5c and 5e). We also observed increases in expression of regulators of triglyceride metabolism, including *Angptl4* and *Angptl8* ^27^, the glycolytic/gluconeogenic enzyme *AldoB*, which is highly upregulated in models of β-cell mitochondrial dysfunction and in T2D islets ^28, 29^, as well as *Nrf1*, a key regulator of mitochondrial biogenesis (Figures 5c, 5e, and S4d; ^30^). To resolve additional mitochondrial specific effectors that were differentially regulated following the loss of Mfn1 and 2, we next overlaid our RNAseq data on MitoCarta 2.0 ^31^, which contains a compendium of targets with strong evidence for localization to the mitochondria (Figure 5d). This revealed increased expression of several genes related to mtDNA replication machinery or associated with the mitochondrial nucleoid (Figures 5d-e), including the critical mtDNA topoisomerase *Top1mt* ^32^, the mitochondrial co-chaperone *Dnaja3*/*Tid1*, which maintains mtDNA integrity ^33^, and the mitochondrial pyrimidine transporter *Slc25a33*, which is essential for mtDNA replication ^34^. We also observed upregulation in the mitochondrial AAA+ protease *LonP1*, which interacts with and regulates the stability of numerous proteins at the mitochondrial nucleoid (Figures 5d-e; ^35–37^). Importantly, we did not observe changes in expression of other fission/fusion genes or regulators of mtRNA transcription, mtDNA repair, sirtuins, or mitophagy (Figure S4d). Taken together, these data indicate that Mfn1/2-deficiency induced a transcriptional signature consistent with an activation of mtDNA replication to potentially compensate for β-cell mtDNA depletion.

### Mfn1 and 2 act through Tfam to regulate β-cell mtDNA copy number

The mechanisms underlying regulation of mtDNA by mitochondrial fusion have been controversial, and previous studies implicate several pathways that could reduce mtDNA content, including genome instability/increased mtDNA mutations, enhanced mitophagy, defects in mitochondria-associated ER membrane (MAM) function, or impaired mtDNA replication ^17, 19, 38–41^. As our studies were completed in young mice, which bear infrequent mtDNA mutations ^42^, and we did not observe differences in expression of mtDNA repair enzymes (Figure S4d), we focused on assessments of mitophagy and mtDNA replication to better resolve the significant reduction of mtDNA observed in β-Mfn1/2^DKO^ islets. We first assessed rates of mitophagy following incubation of control and β-Mfn1/2^DKO^ islets with the cell permeable Mtphagy dye, which did not reveal differences in mitophagy between genotypes (Figure S4e). We also did not observe a difference in frequency of autophagosomes bearing mitochondria by TEM, or changes in essential genes in the mitophagy pathway between control and β-Mfn1/2^DKO^ islets (Figure 2b, S2b and S4d). As an initial screen for MAM function, we measured MAM-dependent lipids, including cholesteryl esters and phospholipids ^43, 44^. However, we did not detect differences in total cholesterol, cholesteryl esters, or phosphatidylserine concentrations in isolated islets of control and β-Mfn1/2^DKO^ mice (Figures S4f-g).

To assess regulators of β-cell mtDNA replication, we examined expression of the replisome proteins Ssbp1, Twinkle, and Polg. We also measured expression of Tfam, a master regulator of mtDNA copy number that promotes mitochondrial genome abundance via effects on mtDNA packaging, stability, and replication, and whose levels are proportional to mtDNA copy number ^11, 45–48^. While replisome proteins were largely unchanged, we observed a significant reduction in Tfam protein in β-Mfn1/2^DKO^ islets (Figures 6a-b). *Tfam* mRNA expression was unchanged in β-Mfn1/2^DKO^ mice (Figure S4d), suggesting the reduction in Tfam protein occurred post-transcriptionally. We further observed a significant increase in the level of LonP1 protein, which has been shown to lead to Tfam protein turnover ^49, 50^. We additionally confirmed reductions in Tfam and increases in LonP1 protein in β-Mfn1^+/-^Mfn2^KO^ islets (Figure S5), correlating with the mtDNA depletion, glucose intolerance, and impaired GSIS observed in these mice (Figures 4c-e). In contrast, β-Mfn1^KO^Mfn2^+/-^ mice, which develop mitochondrial structural alterations without glucose intolerance or mtDNA depletion, did not exhibit changes in Tfam and LonP1 (Figure S5 and Figures 4c-e). These results suggest that changes in Tfam and LonP1 expression following Mfn1/2 deficiency were not due to increases in punctate mitochondria.

**Figure 6.**
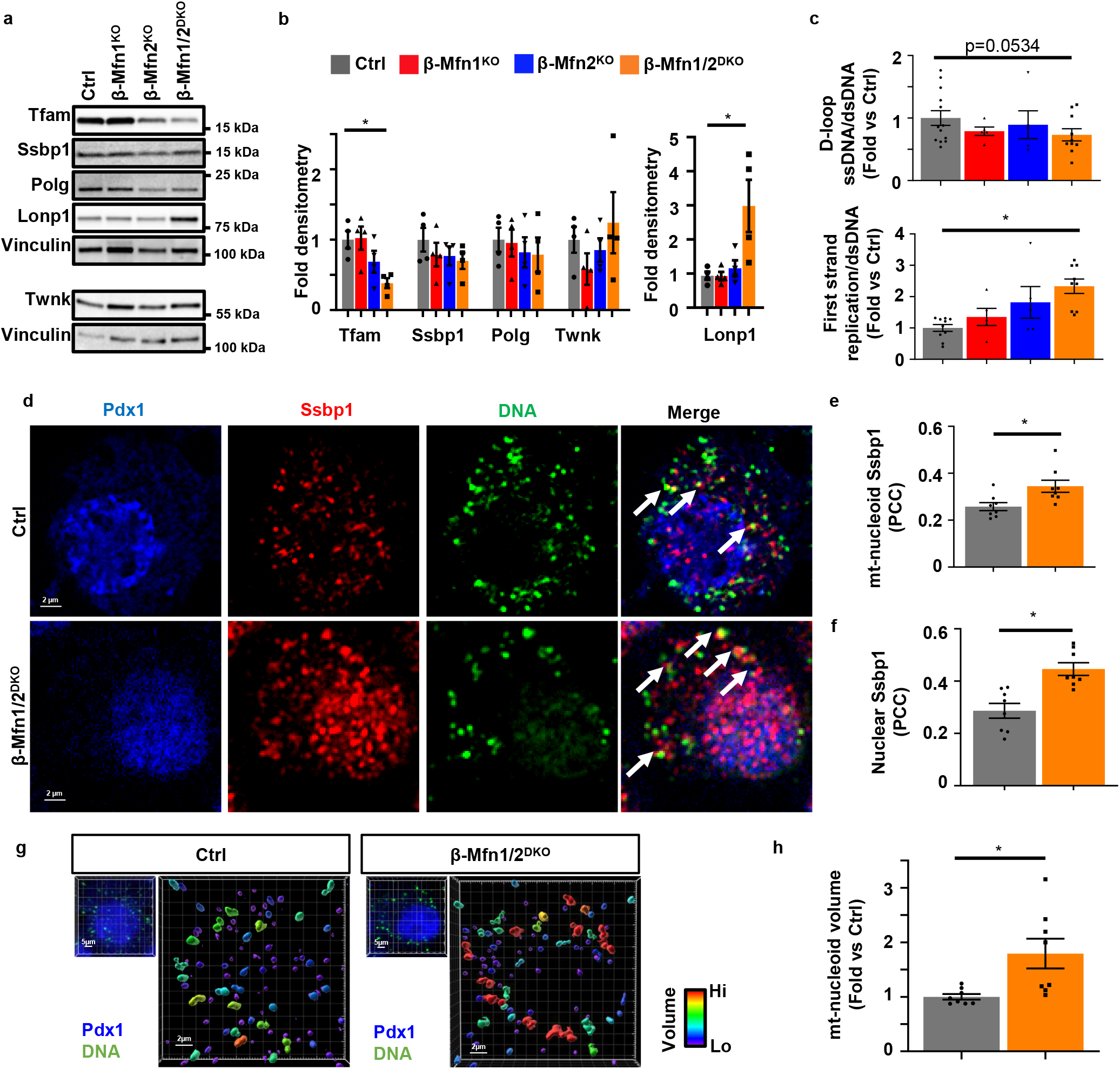
Mfn1 and 2 maintain expression of Tfam in β-cells. **a** Expression of Tfam, key replisome proteins, and LonP1 by Western blot (WB) in islets isolated from 11-week-old Ctrl, β-Mfn1^KO^, β-Mfn2^KO^ and β-Mfn1/2^DKO^ mice. Representative of 4 independent mice per group. **b** Densitometry (normalized to Vinculin) from studies in Figure 6A. n=4/group; Data are presented as mean ± SEM. *p=0.03 for Tfam, p=0.01 for Lonp1 by one-way ANOVA followed by Sidak’s post-hoc test for multiple comparisons. **c** Quantification of single strand mtDNA products measured by qPCR in isolated islets from 8-week-old Ctrl (n=13), β-Mfn1^KO^ (n=5), β-Mfn2^KO^ (n=5) and β-Mfn1/2^DKO^ (n=9) mice. Data are presented as mean ± SEM. *p=0.0007 by one-way ANOVA followed by Sidak’s post-hoc test for multiple comparisons. **d** Deconvolution immunofluorescence image at 100X magnification of islets from Ctrl and β-Mfn1/2^DKO^ mice stained for Ssbp1 (red), mtDNA (green), and Pdx1 (blue). Representative of 8 independent experiments. White arrows demarcate co-localized Ssbp1+ mtDNA+ structures. **e** Quantification of mitochondrial nucleoid Ssbp1 localization (Ssbp1+ mtDNA+ co-localization) in Ctrl and β-Mfn1/2^DKO^ β-cells from studies depicted in Figure 6D by Pearson’s correlation coefficient (PCC). n=8/group. Data are presented as mean ± SEM. *p=0.0142 by two-tailed t-test. (∼100 β-cells from each animal per group were analyzed). **f** Quantification of nuclear Ssbp1 localization in Ctrl and β-Mfn1/2^DKO^ β-cells from studies depicted in Figure 6D by Pearson’s correlation coefficient (PCC). n=8/group. Data are presented as mean ± SEM. *p=0.0008 by two-tailed t-test. (∼100 β-cells from each animal per group were analyzed). **g** Imaris® generated 3D reconstruction of deconvolution immunofluorescence Z-stack images at 100X magnification stained for DNA of β-cells (see inset image – Pdx1, blue, DNA, green) from Ctrl and β-Mfn1/2^DKO^ mice. Colors represent relative mitochondrial volume. Representative image of 8 independent mice/group. **h** Quantification of relative nucleoid volume of Ctrl and β-Mfn1/2^DKO^ β-cells from studies depicted in Figure 6g. n=8/group. Data are presented as mean ± SEM. *p=0.0127 by two-tailed t-test. (∼100 β-cells from each animal per group were analyzed). Uncropped western blots and source data are provided as a Source Data file.

To test for defects in mtDNA replication, we next assessed initiation of first strand replication. First strand synthesis in mitochondrial replication begins in the D-loop. However, a single stranded (ss) molecule that forms the classic D-loop structure called 7S, which is a non-replicative structure, shares this sequence. It is the elongation of the 7S sequence into the nearby region (Cytb) that demonstrates committed initiation of replication. Here we assessed the levels of ss and dsDNA at specific sequences by quantitative PCR to overcome expected sample limitations from studies with mouse pancreatic islets. Notably, import of nascent Tfam has been shown to increase the generation of 7S DNA due to its transcription initiation activity^51^. We found a trend towards reduced total ss 7S sequences normalized to ds mtDNA in β-Mfn1/2^DKO^ islets (Figure 6c). However, the rate of 7S sequence extension into committed first strand replication was increased in β-Mfn1/2^DKO^ islets (Figure 6c), which could represent a compensatory response to mtDNA depletion.

Increased first strand replication intermediates heighten the levels of displaced second strand template, which is bound by the single strand binding protein Ssbp1 ^52^. Consistent with ss qPCR studies, in situ levels of Ssbp1 at the mtDNA nucleoid (detected by perinuclear anti-DNA antibody staining), were significantly elevated in Mfn1/2-deficient β-cells (Figures 6d-e). The increase in Ssbp1 localization to mitochondrial nucleoids could represent binding to a higher quantity of single-stranded mtDNA replication intermediates and could also be consistent with observations of impaired completion of mtDNA replication previously reported following Mfn1/2-deficiency ^19, 53^. We also observed Ssbp1 nuclear localization (Figures 6d and f), which has been previously demonstrated following mitochondrial stress ^54^. Further, we detected an increase in the volume of mtDNA nucleoid structures in Mfn1/2-deficient β-cells by high-resolution 3D imaging (Figures 6g-h), which are similar to previous reports of Tfam-deficiency but could also represent impaired nucleoid distribution ^19, 55, 56^. Taken together, our observations of reduced mtDNA content, mtRNA levels, as well as changes in single-stranded mtDNA products, Ssbp1 localization, and nucleoid size following Mfn1/2-deficiency suggest a role for Tfam in mediating the effects of Mfn1/2 in β-cell function.

To test whether increasing mtDNA copy number was sufficient to improve β-cell function in fusion-deficient islets, we overexpressed Tfam in β-Mfn1/2^DKO^ and littermate control islets. We transduced intact islets using adenoviral vectors encoding the human form of TFAM, thus allowing us to track overexpression of TFAM using human-specific TFAM anti-sera (Figures 7a-b and S6a). Transduction efficiency in β-cells was estimated at ∼40-50% (Figure S6a). Despite limitations in transduction efficiency and expression, TFAM overexpression significantly ameliorated the reductions in both GSIS and mtDNA copy number in β-Mfn1/2^DKO^ islets (Figures 7c-d and S6b). Therefore, Mfn1 and 2 direct β-cell function, at least in part, through the regulation of Tfam-mediated mtDNA copy number control.

**Figure 7.**
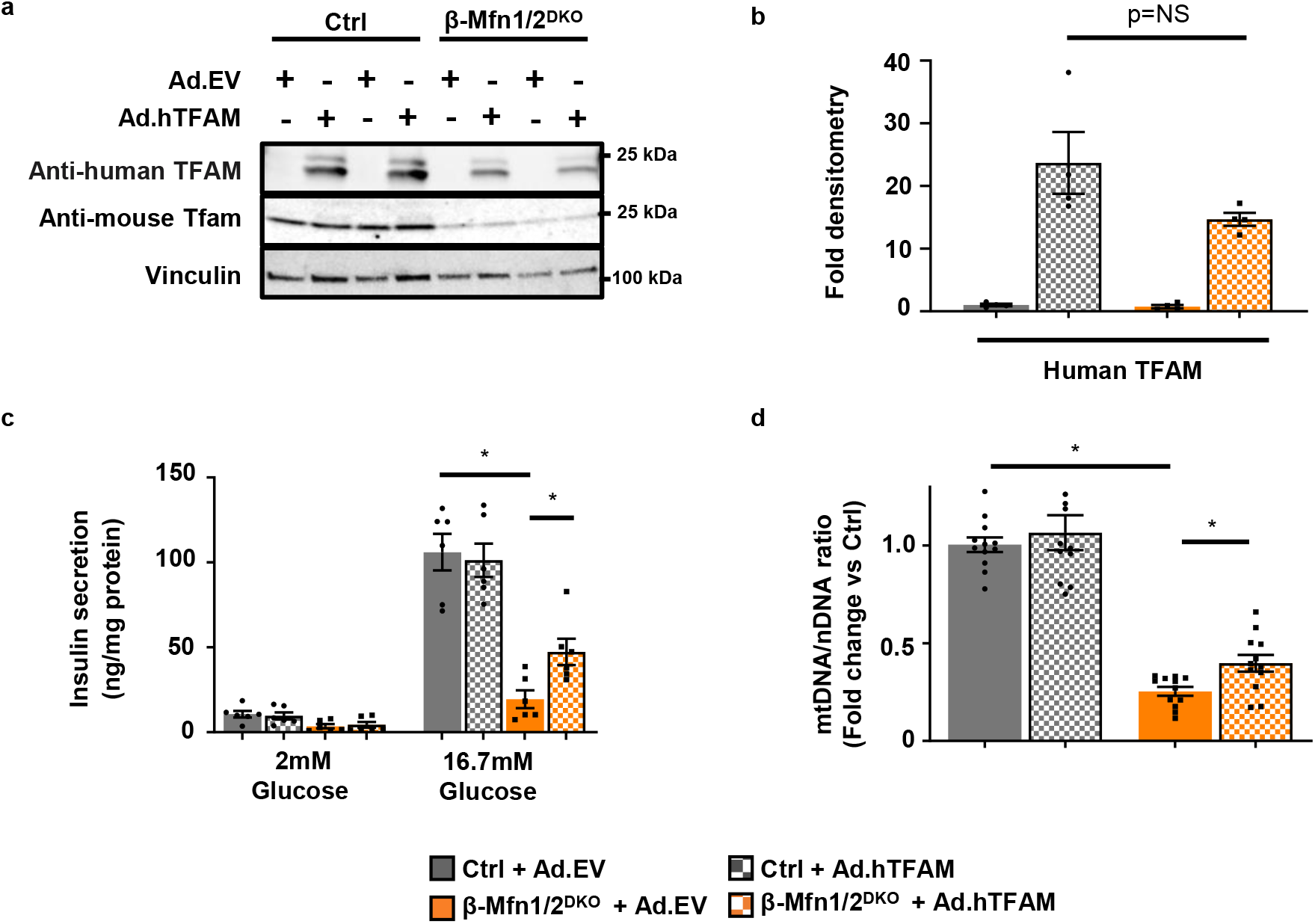
Mfn1 and 2 maintain β-cell function through Tfam-mediated mtDNA copy number control. **a** Expression of mouse-specific Tfam and human-specific TFAM by WB in islets isolated from 10-15-week-old Ctrl and β-Mfn1/2^DKO^ mice, transduced with empty vector control (Ad.EV) or human TFAM-overexpressing (Ad.hTFAM) adenoviral particles. Representative of 4 independent mice/group. **b** Densitometry (normalized to cyclophilin B) from studies in Figure 7A. n=4/group; Data are presented as mean ± SEM. p=NS (non-significant) by two-tailed t-test. **c** Glucose-stimulated insulin secretion following static incubations in 2mM and 16.7 mM glucose, performed in isolated Ctrl and β-Mfn1/2^DKO^ islets following transduction with Ad.EV or Ad.hTFAM adenoviral particles. n=6/group. Data are presented as mean ± SEM. *p<0.0005 for Ctrl + Ad.EV vs β-Mfn1/2^DKO^ + Ad.EV, p=0.0148 for β-Mfn1/2^DKO^ + Ad.EV vs β-Mfn1/2^DKO^ + Ad.hTFAM by one-way ANOVA followed by Tukey’s post-hoc test for multiple comparisons. **d** Relative mtDNA content measured by qPCR (normalized to nuclear DNA expression) from isolated Ctrl and β-Mfn1/2^DKO^ islets following transduction with Ad.EV or Ad.hTFAM adenoviral particles. n=12/group. Data are presented as mean ± SEM. *p<0.0005 for Ctrl + Ad.EV vs β-Mfn1/2^DKO^ + Ad.EV, p=0.007 for β-Mfn1/2^DKO^ + Ad.EV vs β-Mfn1/2^DKO^ + Ad.hTFAM by two-tailed t-test. Uncropped western blots and source data are provided as a Source Data file.

### Mitofusin agonists restore mtDNA content independent of changes in mitochondrial structure

We next asked if pharmacologically targeting mitofusins could restore mtDNA levels in models of mitofusin deficiency independent of changes in mitochondrial structure. To this end, we employed recently described tandem pharmacological agonists, (2-{2-[(5-cyclopropyl-4-phenyl-4H-1,2,4-triazol-3-yl)sulfanyl]propanamido}-4H,5H,6H-cyclopenta[b]thiophene-3-carboxamide and 1-[2-(benzylsulf-anyl)ethyl]-3-(2-methylcyclohexyl)urea (hereafter called Mfn agonists), which together have been previously shown to improve mitochondrial morphology in either Mfn1- or Mfn2-deficient fibroblasts ^57^. These agonists require some residual mitofusin levels to be effective ^57^. Thus, we initially treated control and β-Mfn2^KO^ islets, which exhibit reduced mtDNA content without changes in mitochondrial structure (Figures 3a, 2b-c, and 2e). To avoid systemic effects of modifying mitochondrial fission/fusion balance in other metabolic tissues ^1^, we utilized Mfn agonists on isolated islets. Interestingly, Mfn agonists restored mtDNA content to baseline in β-Mfn2^KO^ islets without affecting mitochondrial morphology or inducing transcriptional regulators of mitochondrial biogenesis (Figures 8a and S7a-b). As anticipated, Mfn agonists were ineffective in the setting of combined absence of Mfn1 and 2 and were unable to rescue reductions in mtDNA content or Tfam protein in β-Mfn1/2^DKO^ islets (Figures S7c-d).

**Figure 8.**
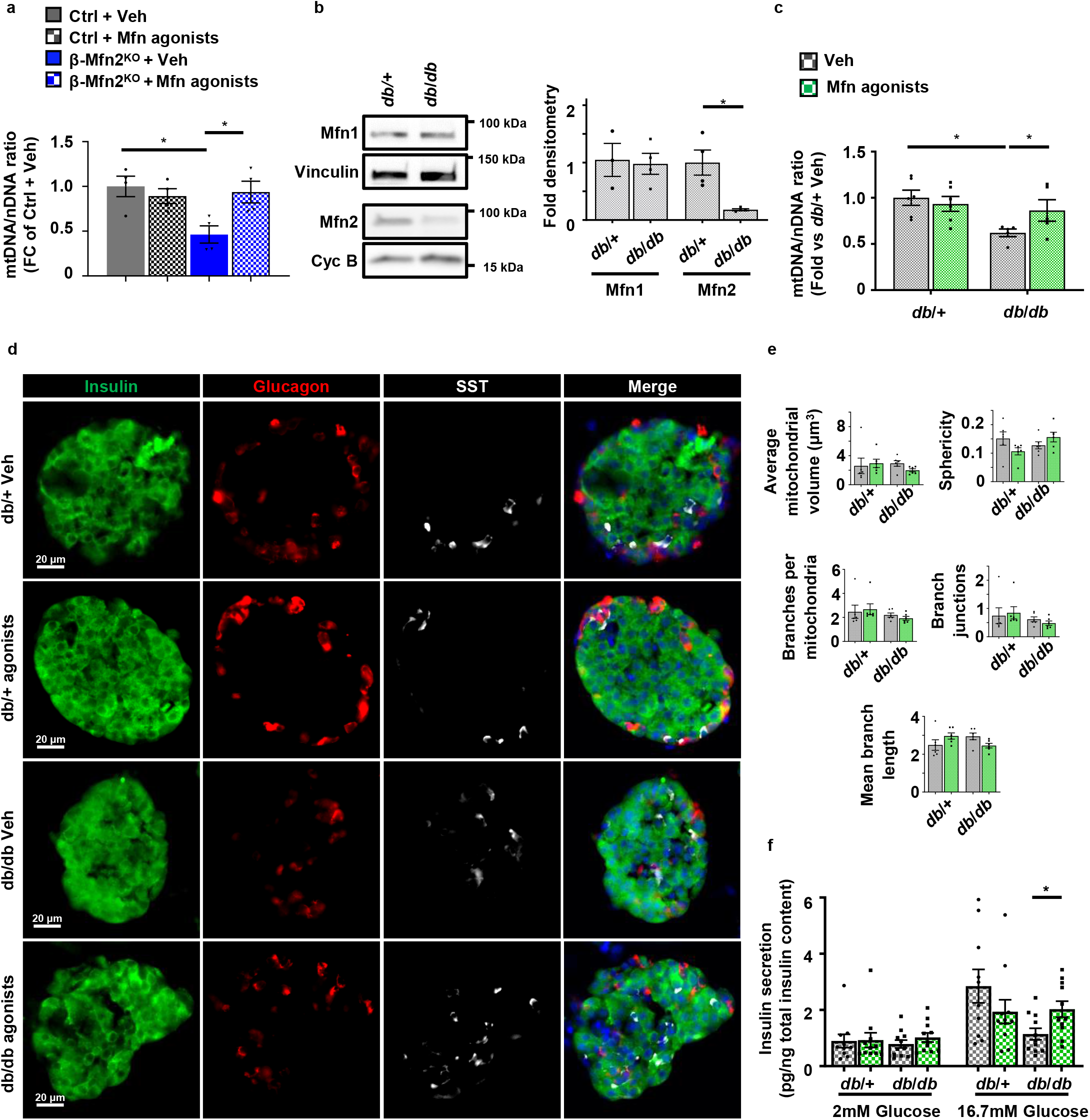
Mitofusin agonists restore mtDNA depletion following Mfn2 deficiency independent of changes in mitochondrial structure. **a** Relative mtDNA content measured by qPCR (normalized to nuclear DNA expression) from isolated islets of Ctrl and β-Mfn2^KO^ mice following treatment with vehicle (Veh) or 0.5 µM Mfn agonists for 24 hrs. n=4/group. Data are presented as mean ± SEM. *p=0.0138 for Ctrl + Veh vs β-Mfn2KO + Veh, p=0.0302 for β-Mfn2^KO^ + Veh vs β-Mfn2^KO^ + Mfn agonists by one-way ANOVA followed by Sidak’s post-hoc test for multiple comparisons. **b** Expression of Mfn1 and Mfn2 by WB in islets isolated from 10-12-week-old *db*/+ and *db*/*db* mice. Vinculin and cyclophilin B serve as loading controls. Representative image (left) of 3 independent mice/group for Mfn1 and 4 independent mice/group for Mfn2. Mfn1 (n=3) and Mfn2 (n=4) densitometry, normalized to vinculin or cyclophilin B (right). Data are presented as mean ± SEM. *p=0.0095 by two-tailed t-test. **c** Relative mtDNA content measured by qPCR (normalized to nuclear DNA expression) from isolated islets of 10-12-week-old *db*/+ (n=6) and *db*/*db* (n=5) mice following treatment with vehicle (Veh) or 0.5 µM mitofusin agonists for 24 hrs. Data are presented as mean ± SEM. *p=0.0038 for *db*/+ + Veh vs *db*/*db* + Veh, p=0.0428 for *db*/*db* + Veh vs *db*/*db* + Mfn agonists by two-tailed t-test. **d** Immunofluorescence images at 20X magnification of islets from 10-12-week-old *db*/+ and *db*/*db* mice stained for Insulin (green), Glucagon (red), Somatostatin (SST; white), and DAPI (blue) following treatment with vehicle (Veh) or 0.5 µM mitofusin agonists for 24 hrs. n=3/group. **e** β-cell mitochondrial morphology and network analysis by MitoAnalyzer of deconvolution immunofluorescence Z-stack images, stained for SDHA (and insulin), from pancreatic islets of 10-12-week-old *db*/+ and *db*/*db* mice following treatment with vehicle (Veh) or 0.5 µM mitofusin agonists for 24 hrs. n=6/group (∼150-200 β-cells/animal were quantified). Data are presented as mean ± SEM. **f** Glucose-stimulated insulin secretion following static incubations in 2 mM and 16.7 mM glucose, performed in isolated islets of 10-12-week-old *db*/+ and *db*/*db* mice following treatment with vehicle (Veh) or 0.5 µM mitofusin agonists for 24 hrs. n=11/group. Data are presented as mean ± SEM. *p=0.0092 by two-tailed t-test. Uncropped western blots and source data are provided as a Source Data file.

We next evaluated the effects of Mfn agonists in other models of mitofusin deficiency, as Mfn2 expression is reduced in several models of β-cell dysfunction ^9, 10^. We assessed the expression of Mfn1 and 2 in islets isolated from the leptin receptor deficient *db*/*db* BKS mouse model, which develops obesity, progressive β-cell mitochondrial and insulin secretory dysfunction, and eventual β-cell failure ^58–62^. We observed that Mfn2, but not Mfn1, protein levels were reduced in islets of 10–12-week-old *db*/*db* mice when compared to *db*/+ controls (Figure 8b), measured at an age prior to β-cell failure ^58, 59^. Similar to observations in β-Mfn2^KO^ islets, *db*/*db* islets exhibited reduced mtDNA content without alterations in mitochondrial structure (Figures 8c and 8e). Concordantly, treatment with Mfn agonists rescued mtDNA levels in *db*/*db* islets, while leaving mitochondrial structure and islet architecture unchanged (Figures 8c-e). We also observed that Mfn agonists improved GSIS in the islets of *db*/*db* mice (Figure 8f). Taken together, these results indicate that Mfn agonists restore mtDNA content and improve GSIS independent of changes in mitochondrial structure. Moreover, these results extend the association between mitofusin deficiency, mtDNA depletion, and impaired GSIS in β-cells.

## DISCUSSION

Here, we identify that Mfn1 and 2 promote mitochondrial fitness to fuel glucose-stimulated insulin release in pancreatic β-cells and maintain glucose homeostasis. Mfn1 or Mfn2 are individually dispensable for β-cell function, but together coordinately control mitochondrial respiration, structure, and mtDNA content. Mfn1 and 2 were thought to regulate cellular metabolism principally through the control of mitochondrial architecture. Our single allele mouse models directly demonstrate that Mfn1/2-deficiency is coupled to glucose intolerance through mtDNA loss rather than mitochondrial structure. Further, we observe that increased mtDNA copy number following Tfam overexpression is capable of ameliorating impaired GSIS in fusion-deficient β-cells. Thus, our studies position the maintenance of mtDNA content as the principal task necessary for Mfn1 and 2 to regulate β-cell function.

Our description of a central role for Mfn1 and 2 in β-cells through mtDNA content rather than mitochondrial structure elucidates a previously unappreciated role for mitochondrial fusion on cellular metabolic function. Mfn1 and 2 act in a complementary manner to support mitochondrial metabolism from development to adulthood. Both germline and cardiomyocyte-specific deletion of Mfn1 and 2 are associated with developmental or early postnatal demise ^16, 18^, presumed to be primarily caused by defects in mitochondrial network dynamics. Later descriptions of reduced mtDNA levels in constitutive or inducible loss of Mfn1/2 in skeletal muscle did not discern the physiologic importance of mtDNA depletion versus structure following fusion-deficiency ^15, 17^. Our results clarify a primary physiologic impact of mtDNA depletion rather than mitochondrial structure following Mfn1/2 deficiency in β-cells that may also underlie the physiologic importance of Mfn1 and 2 in other metabolic tissues. To our knowledge, this is the first time that these functions of Mfn1/2 have been separated in a physiological context.

We observe that β-Mfn1/2^DKO^ islets attempt unsuccessfully to compensate for mtDNA depletion. Our analyses of first strand replication suggest that Mfn1/2-deficient β-cells increase the activation of early steps in mtDNA replication, which is supported by increased Ssbp1 localization to nucleoids to potentially bind newly formed ss-mtDNA. Despite these changes, mtDNA content remains low in Mfn1/2-deficient β-cells, suggesting that the increased commitment to replication is insufficient. Indeed, increased nucleoid Ssbp1 may be engaged in binding abnormal single-stranded replication products due to fusion and replication defects ^19, 53^. The increased expression of *Top1mt* found in Mfn1/2-deficient β-cells (Figure 5e) could also represent a response to increased mitochondrial replication stress ^63^.

Loss of Mfn1/2 in β-cells leads to severe hyperglycemia, reduced GSIS, and mtDNA depletion, which bears similarity to a previous report of selective Tfam deficiency in β-cells ^25^. Notably, previously described β-cell specific Tfam knockout mice developed progressive loss of β-cell mass with age ^25^. We found that β-cell mass was unchanged in young β-Mfn1/2^DKO^ mice, but it is possible that they would similarly develop glucotoxicity and an age-related loss of β-cell mass over time. Alternatively, complete depletion of mtDNA, which occurs in the setting of Tfam-deficiency but does not occur in β-Mfn1/2^DKO^ mice, might be required for loss of β-cell mass. Furthermore, the transgenic *RIP2*-Cre used in the β-cell Tfam knockout model (in contrast to the highly specific *Ins1*-Cre knockin strain we used in β-Mfn1/2^DKO^ mice) is known to result in off-target recombination^25, 64^, which may in part underlie phenotypic differences between the models. In future experiments, it will be intriguing to examine Tfam and Mfn1/2-deficient mice generated using the same Cre-recombinase strains to determine the contribution of Tfam-dependent and independent functions on Mfn1/2 in β-cell health and function.

A related concern in the Mfn1/2-deficiency models is the use of a constitutive Cre strain, which may yield phenotypic defects due to recombination during β-cell development. However, an independent group found that mice with inducible Mfn1/2-deficiency only in adult β-cells developed a highly similar phenotype to constitutive β-Mfn1/2^DKO^ mice ^65^, suggesting that developmental defects are unlikely to be a major contributor to the phenotype we observe.

While our data suggest that Tfam mediates the effects of Mfn1/2 on mtDNA content and GSIS in β-cells, the mechanisms for post-transcriptional loss of Tfam in β-Mfn1/2^DKO^ islets are unclear. Loss of Tfam may be due to increased LonP1 expression, which is known to degrade Tfam ^49, 50^. Further support for a role for LonP1 as a contributor to cellular dysfunction include studies demonstrating reduction of nuclear-encoded subunits of Complex I and III following increases in LonP1 expression ^66–68^, an effect we also observe following Mfn1/2-deficiency (Figure 3d). Mfn1/2-deficiency has also been linked to reductions in nuclear-encoded Complex I and III subunits via instability in respiratory supercomplexes, which have been observed following LonP1 overexpression ^15, 66–68^. However, the function and transcriptional regulation of *LonP1* in β-cells are unknown and require future study.

Mfn2 is frequently observed at MAMs ^69, 70^. MAMs are essential for communication and Ca^2+^ transfer between the ER and mitochondria, are important for β-cell function and GSIS, and have been suggested to regulate mtDNA replication ^39–41, 71–73^. Thus, alterations in MAMs in β-Mfn1/2^DKO^ islets could lead to impaired mtDNA replication. Our initial functional assessments of MAMs by measurement of MAM-dependent lipids did not reveal differences between control and β-Mfn1/2^DKO^ islets (Figures S4f-g). We also did not observe ER dilation or overt changes in mitochondrial-ER proximity upon analysis of transmission EM in β-Mfn1/2^DKO^ islets (Figures 2b and S2b). Additionally, a recent report found that inducible loss of Mfn1/2 in adult β-cells did not alter MAM-dependent ER calcium release ^65^. Despite these findings, our results do not exclude the potential that impaired MAM function at β-cell mtDNA nucleoids contributes to the impairments in mtDNA replication and content we observe. Indeed, our results demonstrating the importance for a single allele of *Mfn2* over *Mfn1* in the maintenance of glycemic control and mtDNA content could suggest a role for Mfn2-dependent MAM function in β-cell health and function, as Mfn1 is not found at MAMs. Alternatively, the regulation of MAMs by Mfn2, as well as the MAM-dependent role of Mfn2 in mtDNA replication, remains controversial ^74, 75^. Recent work also implicates mitochondrial fusion as necessary to maintain the stoichiometry of the protein components within the mtDNA replisome ^19^. Future studies will be valuable to dissect the molecular details underlying the connections between mitochondrial fusion, MAMs, and control of β-cell mtDNA replication.

Mitochondrial structural defects and age-related mtDNA depletion combine to impair β-cell function to increase T2D risk ^6, 7, 76–78^, thus, improving mitochondrial function would have significant translational benefits to restore β-cell health in diabetes. Our studies demonstrate the feasibility of targeting mtDNA copy number to improve GSIS in the setting of mitofusin deficiency. While our results in *db*/*db* islets are promising, loss of the leptin receptor could lead to mitochondrial defects that are particularly amenable to improvement by Mfn agonists. Thus, it will be intriguing to determine if Mfn agonists can also reverse β-cell dysfunction in diet-induced models of obesity *in vivo*. Further, recently identified compounds targeting pyrimidine metabolism to induce *Mfn1/2* transcription ^79^ will be of interest to determine if they can improve β-cell function and mtDNA content. Our observation of the upregulation of mitochondrial pyrimidine transporter *Slc25a33* in β-Mfn1/2^DKO^ islets is consistent with links between pyrimidine metabolism and mitochondrial fusion, which suggest these compounds could hold promise to enhance β-cell mitochondrial function. As expression of MFN1 and MFN2 is reduced in cardiomyocytes and skeletal muscle of patients with diabetes, respectively ^80–82^, activators of Mfn1/2 may have additional benefits beyond improving β-cell function. Future studies will be required to fully characterize the efficacy of overcoming mitochondrial defects *in vivo* to treat diabetes.

## METHODS

### Animals

*Mfn1*^loxP/loxP^ and *Mfn2*^loxP/loxP^ mice possessing loxP sites flanking exon 4 of the *Mfn1* gene and exon 6 of the *Mfn2* gene, respectively, were purchased from Jackson Laboratories ^13^. *Mfn1*^loxP^ and *Mfn2*^loxP^ mice were mated to *Ins1*-Cre ^83^ to generate experimental groups. *Ins1*-Cre–alone, *Mfn1*^loxP/loxP^ and *Mfn2*^loxP/loxP^ mice were phenotypically indistinguishable from each other and combined as controls (Ctrl; see Figures S1b-c). *Ins1*-Cre–alone were also phenotypically indistinguishable from wild-type C57BL/6N controls, consistent with previous reports from our group and others ^21, 83^. All animals were maintained on the C57BL/6N background. *db*/+ and *db*/*db* mice, maintained on the BKS background, were purchased from Jackson Laboratories. Animals were housed on a standard 12-hour light/12-hour dark cycle at room temperature and 30-70% humidity, with ad libitum access to food and water. All mice were fed PicoLab® Laboratory Rodent Diet 5L0D (LabDiet) containing 28% protein, 13% fat, and 57% carbohydrates. All studies and endpoints in all figures were completed using both male and female mice, and results from both sexes were combined in all experimental groups. Animal studies were approved by the University of Michigan Institutional Animal Care and Use Committee.

### Mitochondrial respirometry

Islet respirometry was measured using an XF96 extracellular flux analyzer (Seahorse Bioscience) according to manufacturer’s instructions. Briefly, 20 similarly sized islets were plated per well of a Seahorse spheroid plate pre-treated with CellTak (Corning), similar to previously published approaches ^84^. Prior to respirometry profiling, islets were incubated for 1 hour in an atmospheric CO_2_ incubator at 37°C and supplemented with mouse islet culture media^85, 86^ comprised of pH 7.4 unbuffered RPMI1640 media (Seahorse Bioscience) and 2mM glucose prior to flux analyzer assays. Analysis was performed at baseline, upon exposure to 20mM glucose, and again after treatment with 2 µM antimycin A. OCR measurements were normalized to islet number confirmed by light microscopy (Zeiss Stemi 2000) after the completion of respirometry assays.

### Transmission electron microscopy

Mouse islets were pelleted, fixed in 3% glutaraldehyde and 3% paraformaldehyde in 0.1 M Cacodylate buffer (CB; pH 7.2) overnight at 4°C. Islets were then embedded in 2% agarose as described previously ^87^. Samples were then subjected to osmification in 1.5% K_4_Fe(CN)_6_ + 2% OsO_4_ in 0.1 CB for 1 h, dehydrated by serial washes in EtOH (30%, 50%, 70%, 80%, 90%, 95% and 100%) and embedded in Spurr’s resin by polymerization at 60° C for 24h. Polymerized resins were then sectioned at 90nm thickness using a Leica EM UC7 ultramicrotome and imaged at 70kV using a JEOL 1400 TEM equipped with an AMT CMOS imaging system. Mitochondrial structures (Aspect Ratio, Perimeter, Circularity) were analyzed and quantified by ImageJ. Form factor assessments were analyzed by ImageJ and quantified as described ^22^.

### Imaging of islet and organelle morphology and subcellular localization analysis

Mitochondrial morphology and subcellular localization analyses were performed on immuno-stained paraffin-embedded mouse pancreas tissue sections and dissociated islet cells as previously described ^21^. Briefly, pancreatic tissue sections were fixed in 4% PFA overnight, embedded in paraffin and sectioned. Following dewaxing and rehydration, antigen retrieval was performed with 10mM citric acid (pH 6.0). Dissociated islet cells were fixed in 4% PFA for 15 minutes and centrifuged onto glass slides. Pancreatic sections and dissociated islets were then blocked using 5% donkey serum for 1hr at room temperature followed by immunostaining. Islet architecture was analyzed using isolated mouse islets fixed in 4% PFA overnight followed by a 20 min incubation with methylene-blue solution (Sigma) and OCT embedding. Cryo-sectioned islets were rinsed in 75% ethanol for 5 minutes to remove methylene blue, permeabilized with 0.2% Triton-X 100 for 15 minutes, blocked using 5% donkey serum for 1h and immunostained.

Images were captured with an IX81 microscope (Olympus) using an ORCA Flash4 CMOS digital camera (Hamamatsu). Immunostained pancreatic sections, cryo-sectioned and dispersed mouse islets were captured with Z-stack images and subjected to deconvolution (CellSens; Olympus). Mitochondrial morphology and nucleoids were visualized using 3D-renderings generated with Imaris® imaging software (Bitplane). Quantitative 3D assessments of mitochondrial morphology and network were performed on ImageJ using Mitochondria Analyzer plugin ^22^. Co-localization analyses were performed on Z-stack images of immunostained dissociated islet cells using the Coloc2 plugin on ImageJ. Changes in relative nucleoid size were quantified from 3D deconvolution images of immuno-stained dissociated islet cells, detected by perinuclear anti-DNA antibody staining, using ImageJ.

### mtDNA content and replication assays

Relative mtDNA content by qPCR with SYBR-based detection (Universal SYBR Green Supermix; Biorad) was conducted as previously described ^88^ following DNA isolation with the Blood/Tissue DNeasy kit (Qiagen) according to manufacturer’s protocols. First strand mtDNA replication levels were measured by qPCR following digestion by Mnl I, which site specifically digest double stranded DNA ^89^. Briefly, primers specific to 7S mtDNA (5’-ACTCTTCTCTTCCATATGACTATCCC-3’ and 5’-GGCCCTGAAGTAAGAACCAGATGT-3’), whose amplicon contains a Mnl I site, were used to quantify relative single-stranded mtDNA forming the D-loop. Single-stranded DNA generated from the extension of 7S region was measured using primers specific to CYTB (5’-TGCATACGCCATTCTACGCTCA-3’ and 5’-GTGATTGGGCGGAATATTAGGCTTC-3’), which also contains an Mnl I site. Primers specific to COXI (5’-GCAGGAGCATCAGTAGACCTAACA-3’ and 5’-GCGGCTAGCACTGGTAGTGATAAT-3’), whose amplicon does not contain an Mnl I site, were used to quantify non-replicating double stranded mtDNA. The ratio of 7S or CYTB amplicons to the COXI amplicon were then calculated after qPCR of Mnl I digested DNA to determine the relative amount of single stranded mtDNA at the D-loop strand or initiation of first strand replication, respectively.

### RNA sequencing, differential expression, and functional analysis

Sequencing was performed by the Advanced Genomics Core at University of Michigan Medical School. Total RNA was isolated, and DNase-treated using commercially available kits (Omega Biotek and Ambion, respectively). Libraries were constructed and subsequently subjected to 151bp paired-end cycles on the NovaSeq-6000 platform (Illumina). FastQC (v0.11.8) was used to ensure the quality of data. Reads were mapped to the reference genome GRCm38 (ENSEMBL), using STAR (v2.6.1b) and assigned count estimates to genes with RSEM (v1.3.1). Alignment options followed ENCODE standards for RNA-seq. FastQC was used in an additional post-alignment step to ensure that only high-quality data were used for expression quantitation and differential expression. Data were pre-filtered to remove genes with 0 counts in all samples. Differential gene expression analysis was performed using DESeq2, using a negative binomial generalized linear model (thresholds: linear fold change >1.5 or <-1.5, Benjamini-Hochberg FDR (P_adj_) <0.05). Plots were generated using variations of DESeq2 plotting functions and other packages with R version 3.3.3. Genes were annotated with NCBI Entrez GeneIDs and text descriptions. Functional analysis, including candidate pathways activated or inhibited in comparison(s) and GO-term enrichments, was performed using iPathway Guide (Advaita). The RNA sequencing data generated in this study have been deposited in the Gene Expression Omnibus (GEO) database under accession code <u>GSE194204</u>.

### Adenoviral transfection

Isolated mouse islets were distended with Accutase for 1 min at 37° C as described previously^90^. Islets were then transduced with 0.15 MOI of Ad.EV (expressing an empty vector) or Ad.hTFAM (expressing human-specific TFAM) for 48 hrs (viral particles purchased from Vector Biolabs). β-cell transduction was evaluated on dissociated islet cells by immunostaining with Pdx1 and human-specific TFAM anti-sera.

### Flow cytometry

Following isolation and culture, live mouse islets were exposed to 100 nM Mtphagy dye (Dojindo Molecular Technologies) for 3 hours to assess time-dependent accumulation of mitochondria to acidic organelles by the relative fluorescence intensity of the dye per cell as described ^91, 92^. Islets were then dissociated into single cells, stained with DAPI (Thermo Fisher Scientific) and Fluozin-3 (Thermo Fisher Scientific), and resuspended in phenol red–free islet culture medium as previously described ^21^. Samples were analyzed on an LSR Fortessa flow cytometer (BD Biosciences). Single cells were gated using forward scatter and side scatter (FSC and SSC, respectively) plots, DAPI staining was used to exclude dead cells, and Fluozin-3 was used to identify β-cells as previously described ^21, 93^. Mtphagy measurements in β-cells were made using 488 nm excitation laser with a 710 nm emission filter and analyzed using FlowJo (Tree Star Inc.). A total of 5,000 β-cells was quantified from each independent islet preparation. Gating strategies with representative plots are provided in Supplementary Figure S8.

### Glucose and insulin measurements

IPGTT, ITT, and serum insulin measurements were performed as described previously ^85, 86^. Briefly, mice were fasted for 6 hrs prior to glucose and insulin measurements. For IPGTT or ITT, mice were injected intraperitoneally with 2 g/kg glucose or 0.75 U/kg Insulin (Novolin R), respectively. Tail blood was collected for measurements of blood glucose. For *in vivo* insulin release measurements, tail blood was collected following glucose injection, centrifuged at 2000 rpm for 10 min at 4 °C and stored at −20 °C. Static insulin secretion assays on isolated islets were performed following 1hr incubations with 2mM and 16.7mM glucose in HEPES-supplemented KRB buffer containing 135 mM NaCl, 4.7 mM KCl, 1.2 mM KH_2_PO_4_, 5 mM NaHCO_3_, 1.2 mM MgSO_4_.7H_2_O, 1 mM CaCl_2_, 10 mM HEPES and 0.1% BSA (pH 7.4). Samples were analyzed for insulin release using Mouse Ultrasensitive Insulin ELISA (ALPCO) as per the manufacturer’s protocol.

### Islet isolation and cell culture

Mouse islets were isolated and cultured as previously described ^85, 86^. Cell treatments included DMSO (Fisher) and antimycin A (Sigma). Mitofusin agonist treatment included an equimolar mixture of two previously described compounds (0.5 µM each; ^57^), which were administered for 24 hours (2-{2-[(5-cyclopropyl-4-phenyl-4H-1,2,4-triazol-3-yl)sulfanyl]propanamido}-4H,5H,6H-cyclopenta[b]thiophene-3-carboxamide, CAS# 920868-45-7 and 1-[2-(benzylsulfanyl)ethyl]-3-(2-methylcyclohexyl)urea, CAS# 1007818-44-1; Enamine Ltd.).

### Western blotting, quantitative PCR, and immunostaining

All assays were performed as previously described ^21, 85, 86^. Mouse pancreatic islets were lysed with radioimmunoprecipitation assay buffer containing protease and phosphatase inhibitors (Calbiochem), and insoluble material was removed by centrifugation. Equal amounts of proteins were resolved on 4%–20% gradient Tris-glycine gels (Bio-Rad) and transferred to nitrocellulose membranes (Bio-Rad). Membranes were then blocked in 5% milk for 1hr and immunoblotting was performed using Cyclophilin B (1:5000; ThermoFisher, Catalog# PA1-027A), Drp1 (1:1000; CST #8570), Lonp1 (1:1000, ProteinTech, Catalog# 15440-1-AP), Mfn1 (1:1000; Abcam, Catalog# ab126575), Mfn2 (1:1000; Abcam, Catalog# ab56889), mt-Cytb (1:500; ProteinTech, Catalog# 55090-1-AP), Opa1 (1:1000, BD Transduction laboratories, Catalog# 612606), Total Oxphos antibody cocktail (1:1000; Abcam, ab110413), Polg (1:1000, Abcam, Catalog# ab128899), Ssbp1 (1:250; Atlas antibodies; Catalog# HPA002866), Anti-human TFAM (1:1000, PhosphoSolutions, Catalog# 1999-hTFAM), Anti-mouse TFAM (1:1000; PhosphoSolutions 2001-TFAM), Tom20 (1:1000; Cell Signaling Technology, Catalog# 42406), Twnk (1:1000, ProteinTech, Catalog# 13435-1-AP), Vinculin (1:5000; Millipore, Catalog# CP74) and species-specific HRP-conjugated secondary antibodies (Vector Laboratories). Immunostaining on pancreatic sections, cryo-sectioned and dispersed mouse islets was performed using Anti-DNA (1:50; American Research Products, Catalog# PA1-027A), Insulin (1:250; Dako, Catalog# A0564), Insulin (1:100, Abcam, Catalog# ab7842), Glucagon (1:2000; SantaCruz, Catalog# sc-13091), Pdx1 (1:250; Abcam, Catalog# ab47383), SDHA (1:100, Abcam, Catalog# ab14715), Ssbp1 (1:250; Atlas antibodies, Catalog# HPA002866), Somatostatin (1:500, Abcam, Catalog# ab30788) and species-specific Cy2, Cy3 and Cy5-conjugated secondary antibodies (Jackson Immunoresearch). β-cell mass was quantified as total insulin-positive area/total pancreatic area multiplied by pancreatic wet weight using stitched images of complete pancreatic sections ^86^. All antibodies used for Western blotting and immunostaining are listed in Supplementary Table 1.

RNA samples were reverse transcribed using High-Capacity cDNA Reverse Transcription Kit (Thermo Fisher Scientific). Quantitative reverse transcription PCR (qRT-PCR) was performed with SYBR-based detection (Universal SYBR Green Supermix; Biorad) using primers for *Ppargc1a* (5’-ACTATGAATCAAGCCACTACAGAC-3’ and 5’- TTCATCCCTCTTGAGCCTTTCG-3’), *Nrf1* (5’-ACAGATAGTCCTGTCTGGGGAAA-3’ and 5’- TGGTACATGCTCACAGGGATCT-3’) and *Tfam* (5’-AGCTTCCAGGAGGCAAAGGATGAT-3’ and 5’-ACTTCAGCCATCTGCTCTTCCCAA-3’).

### Cholesterol esterification and phosphatidylserine assay

Total amounts of free cholesterol and cholesterol esters were determined in mouse islet lysates as per the manufacturer’s instructions (Promega). Phosphatidylserine levels were measured in mouse islet lysates as per the manufacturer’s instructions (LSBio).

### Statistics

In all figures, data are presented as means ± SEM, and error bars denote SEM, unless otherwise noted in the legends. Statistical comparisons were performed using unpaired two-tailed Student’s t-tests, one-way or two-way ANOVA, followed by Tukey’s or Sidak’s post-hoc test for multiple comparisons, as appropriate (GraphPad Prism). A *P* value < 0.05 was considered significant.

### Data Availability

The authors declare that data supporting the findings of this study are available within the article and its supplementary information files or from the corresponding author (S.A.S.) upon request. RNA sequencing data used for differential gene expression-, pathway- and GO-enrichment analyses have been deposited in the GEO database under the accession number GSE194204. The reference genome used for RNA sequencing was obtained from ENSEMBL (Mus_musculus - Ensembl genome browser 105). All uncropped western blots and data values for all figures are provided in the source data file.

### Code Availability

High-throughput RNA sequencing data was analyzed using DESeq2 (v1.26.0) to determine differential gene expression. iPathway Guide (Advaita version 1910) was used to perform pathway and GO-term analysis.

## Acknowledgements

S.A.S was supported by the JDRF (CDA-2016-189, SRA-2018-539, COE-2019-861), the NIH (R01 DK108921, U01 DK127747), the Department of Veterans Affairs (I01 BX004444), the Brehm family, and the Anthony family. E.L.-D. was supported by the NIH (T32 AI007413 and T32-AG000114). G.L.P. was supported by the American Diabetes Association (19-PDF-063). B.A.K. was supported by the Department of Veterans Affairs (I01 BX004444). The JDRF Career Development Award to S.A.S. is partly supported by the Danish Diabetes Academy and the Novo Nordisk Foundation. We acknowledge the Microscopy, Imaging and Cellular Physiology Core of the University of Michigan DRC (P30 DK020572) for assistance with imaging studies. We thank the University of Michigan Flow Cytometry Core for assistance with flow cytometry studies. Next generation sequencing was carried out in the Advanced Genomics Core at the University of Michigan. We acknowledge support from the Bioinformatics Core of the University of Michigan’s Biomedical Research Core Facilities. We thank Drs. K. Claiborn, C. Rutledge, N. Desai, G. Rutter, A. Garner, A. Menon, M. Torres, and members of the Soleimanpour laboratory for helpful advice.

## Author Contributions

V.S. conceived, designed and performed experiments, interpreted results, drafted and reviewed the manuscript. J.Z. and E.L-D. designed and performed experiments, interpreted results, and reviewed the manuscript. G.L.P. and E.C.R. designed and performed experiments and interpreted results. E.M.W. designed and performed experiments, interpreted results, edited, and reviewed the manuscript. B.A.K. designed studies, interpreted results and reviewed the manuscript. S.A.S. conceived and designed the studies, interpreted results, drafted, edited, and reviewed the manuscript.

