## Supplementary figures for "Mitofusin 1 and 2 regulation of mitochondrial DNA content is a critical determinant of glucose homeostasis"

**Supplementary Table 1.** List of antibodies used within this study.

| Antibody | Company | Catalog # |
|---|---|---|
| Cyclophilin B | ThermoFisher | PA1-027A |
| Anti-DNA | American Research Products (ARP) | 03-61014 |
| Drp1 | Cell Signaling Technology | 8570 |
| Glucagon | SantaCruz | sc-13091 |
| Insulin | Dako | A0564 |
| Insulin | Abcam | ab7842 |
| LonP1 | Proteintech | 15440-1-AP |
| Mfn1 | Abcam | ab126575 |
| Mfn2 | Abcam | ab56889 |
| mt-Cytb | ProteinTech | 55090-1-AP |
| Opa1 | BD Transduction laboratories | 612606 |
| Total OXPHOS antibody cocktail | Abcam | ab110413 |
| Pdx1 | Abcam | ab47383 |
| Polg | Abcam | ab128899 |
| SDHA | Abcam | ab14715 |
| Somatostatin | Abcam | ab30788 |
| Ssbp1 | Atlas antibodies | HPA002866 |
| Anti-human TFAM | PhosphoSolutions | 1999-hTFAM |
| Anti-mouse Tfam | PhosphoSolutions | 2001-TFAM |
| Tom20 | Cell Signaling Technology | 42406 |
| Twnk | Proteintech | 13435-1-AP |
| Vinculin | Millipore | CP74 |

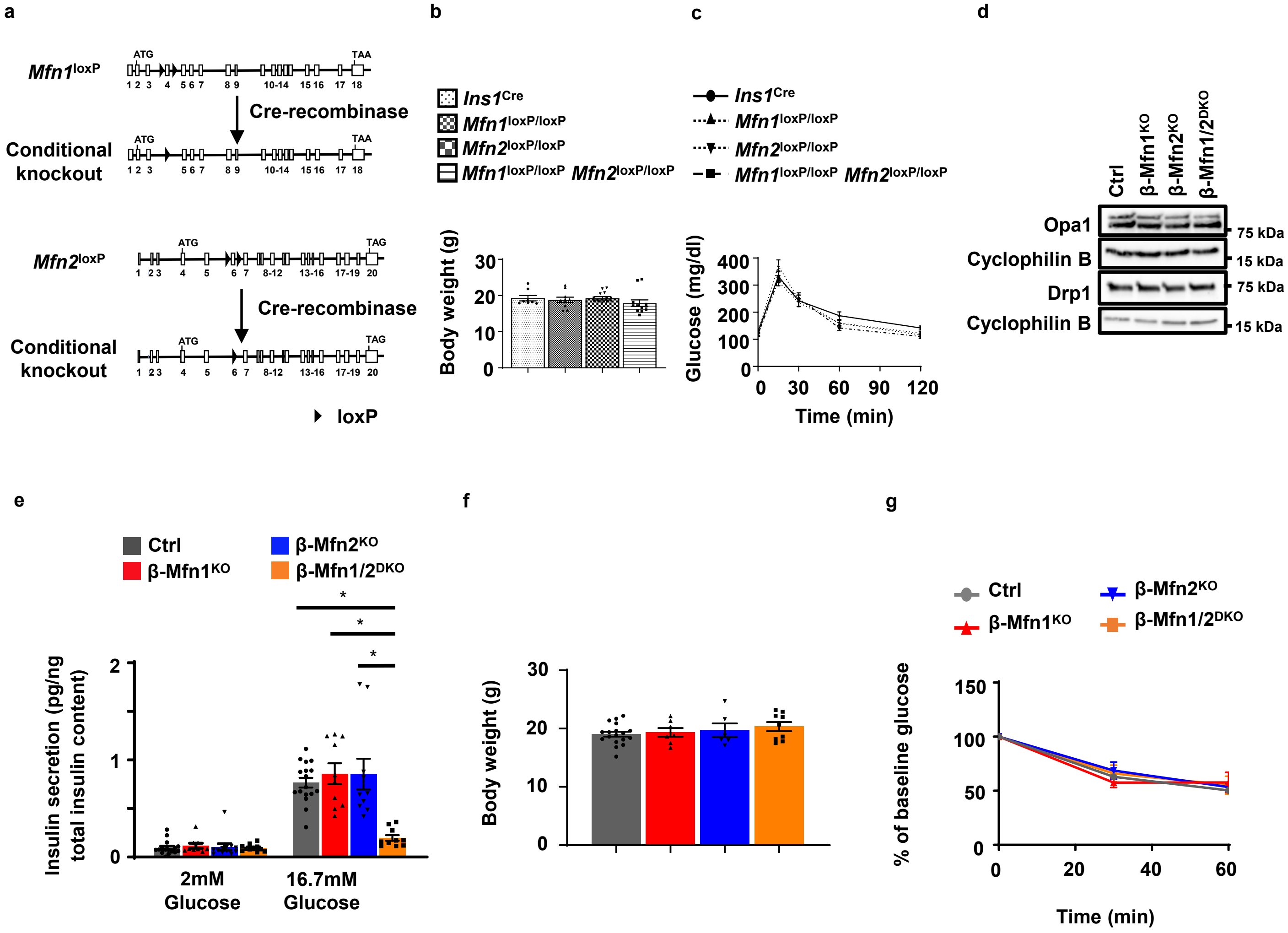


**Figure S1. Body weight and insulin sensitivity are unchanged following loss of Mfn1 and/or Mfn2 in β-cells.**

**a** Schematic representation of conditional targeting of exons 4 and 6 in *Mfn1* (top) and *Mfn2* (bottom), respectively, by Cre-mediated recombination. b Total body weight measured in 8-week-old *Ins1*$^{Cre}$ (n=8), *Mfn1*$^{loxP/loxP}$(n=10), *Mfn2*$^{loxP/loxP}$ (n=14), and *Mfn1*$^{loxP/loxP}$*Mfn2*$^{loxP/loxP}$ (n=12) littermates. Data are presented as mean ± SEM. **c** Blood glucose concentrations measured during IPGTT of 8-week-old *Ins1*$^{Cre}$ (n=8), *Mfn1*$^{loxP/loxP}$ (n=10), *Mfn2*$^{loxP/loxP}$ (n=14), and *Mfn1*$^{loxP/loxP}$*Mfn2*$^{loxP/loxP}$ (n=12) littermates. Data are presented as mean ± SEM. **d** Expression of Opa1 and Drp1 by Western blot (WB) in islets isolated from 8-10-week-old Ctrl, β-Mfn1$^{KO}$, β-Mfn2$^{KO}$ and β-Mfn1/2$^{DKO}$ mice. Vinculin and cyclophilin B serve as loading controls. Representative of 10 independent mice/group for Opa1 and 6 independent mice/group for Drp1. **e** Glucose-stimulated insulin secretion (normalized to total islet insulin content) following static incubations in 2mM and 16.7 mM glucose performed in isolated islets of 8-week-old Ctrl (n=16), β-Mfn1$^{KO}$ (n=8), β-Mfn2$^{KO}$ (n=11), and β-Mfn1/2$^{DKO}$ (n=10) littermates. Data are presented as mean ± SEM. *p<0.0001 by one-way ANOVA followed by Sidak's post-hoc test for multiple comparisons. **f** Total body weight measured in 10-week-old Ctrl (n=18), β-Mfn1$^{KO}$ (n=7), β-Mfn2$^{KO}$ (n=6), and β-Mfn1/2$^{DKO}$ (n=10) mice. Data are presented as mean ± SEM. **g** Blood glucose concentrations (presented as % of baseline glucose) measured during insulin tolerance testing (ITT) of 8-week-old Ctrl (n=19), β-Mfn1$^{KO}$ (n=5), β-Mfn2$^{KO}$ (n=5), and β-Mfn1/2$^{DKO}$ (n=12) littermates. Data are presented as mean ± SEM. Uncropped westerns and raw data are provided as a Source Data file.

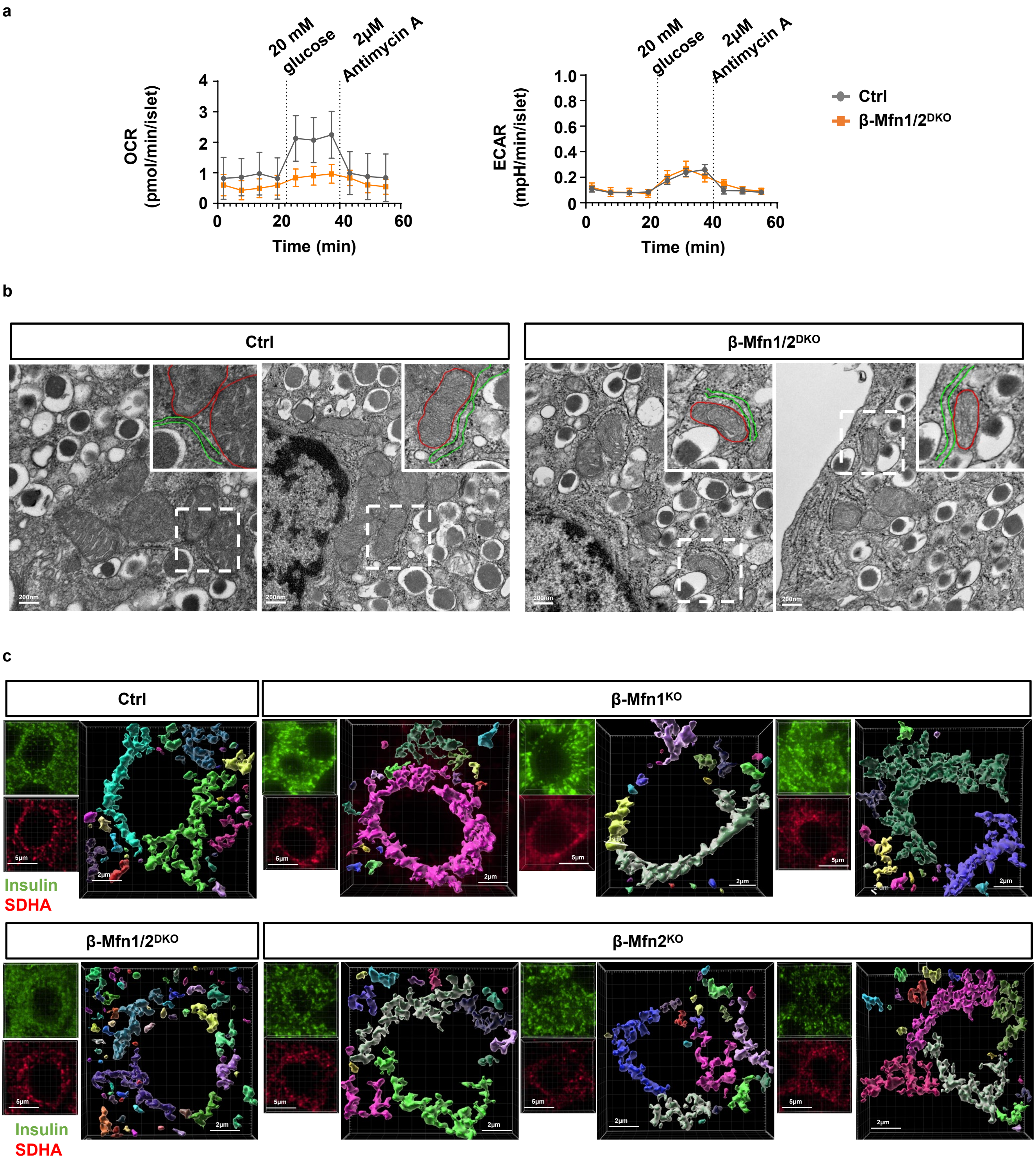


**Figure S2. Evaluation of oxygen consumption rates, mitochondrial/ER ultrastructure, and mitochondrial networking following Mfn1 and/or Mfn2 deficiency in β-cells.**

**a** Representative trace of OCR (left) and ECAR (right) measured in islets isolated from Ctrl and β-Mfn1/2$^{DKO}$ mice following exposure to 2mM glucose, 20mM glucose, and 2μM antimycin A by Seahorse flux analyzer. Data are presented as mean ± SD. **b** Representative transmission EM images of mitochondria and ER (dotted lines delineate region visible in magnified inset, upper right) in β-cells from 12-week-old Ctrl and β-Mfn1/2$^{DKO}$ littermates. Inset - mitochondria and ER are outlined in red and green, respectively. n=3/group. **c** Imaris® generated three-dimensional reconstruction of deconvolution immunofluorescence Z-stack images at 100X magnification stained for SDHA (see inset image – red) from pancreatic sections of β-Mfn1$^{KO}$ and β-Mfn2$^{KO}$ mice (Ctrl and β-Mfn1/2$^{DKO}$ are provided as comparative reference). β-cells were identified by insulin co-staining (inset: insulin – green). Each unique color represents a separate β-cell mitochondrial network cluster. Representative images of 6 independent mice/group. Source data are provided as a Source Data file.

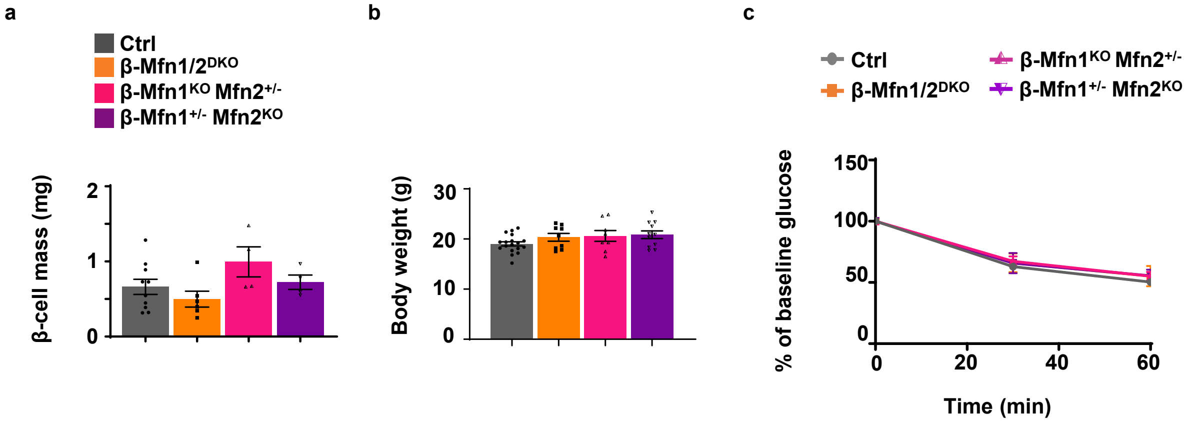


**Figure S3. Defects in β-cell mass, body weight, or insulin sensitivity are not observed in mice bearing only a single allele of *Mfn1* or *Mfn2* in β-cells.**

**a** Pancreatic β-cell mass (n=4-10/group) measured in 10-week-old Ctrl (n=10), β-Mfn1/2$^{DKO}$ (n=6), β-Mfn1$^{KO}$Mfn2$^{+/-}$ (n=4), and β-Mfn1$^{+/-}$Mfn2$^{KO}$ (n=4) littermates. Data are presented as mean ± SEM. **b** Total body weight measured in 10-week-old Ctrl (n=18), β-Mfn1/2$^{DKO}$ (n=10), β-Mfn1$^{KO}$Mfn2$^{+/-}$ (n=8), and β-Mfn1$^{+/-}$Mfn2$^{KO}$ (n=11) littermates. Data are presented as mean ± SEM. **c** Blood glucose concentrations (presented as % of baseline glucose) measured during insulin tolerance testing (ITT) of 8-week-old Ctrl (n=19), β-Mfn1/2$^{DKO}$ (n=12), β-Mfn1$^{+/-}$Mfn2$^{KO}$ (n=13), and β-Mfn1$^{KO}$Mfn2$^{+/-}$ (n=12) littermates. Data are presented as mean ± SEM. Of note, studies in Ctrl and β-Mfn1/2$^{DKO}$ mice were performed together alongside all β-Mfn1$^{KO}$, β-Mfn2$^{KO}$, β-Mfn1$^{+/-}$Mfn2$^{KO}$, and β-Mfn1$^{KO}$Mfn2$^{+/-}$ littermates and thus appear twice for purposes of relevant comparisons (in Figures 1F, S1F, S1G and again in Figures S3A-C). Source data are provided as a Source Data file.

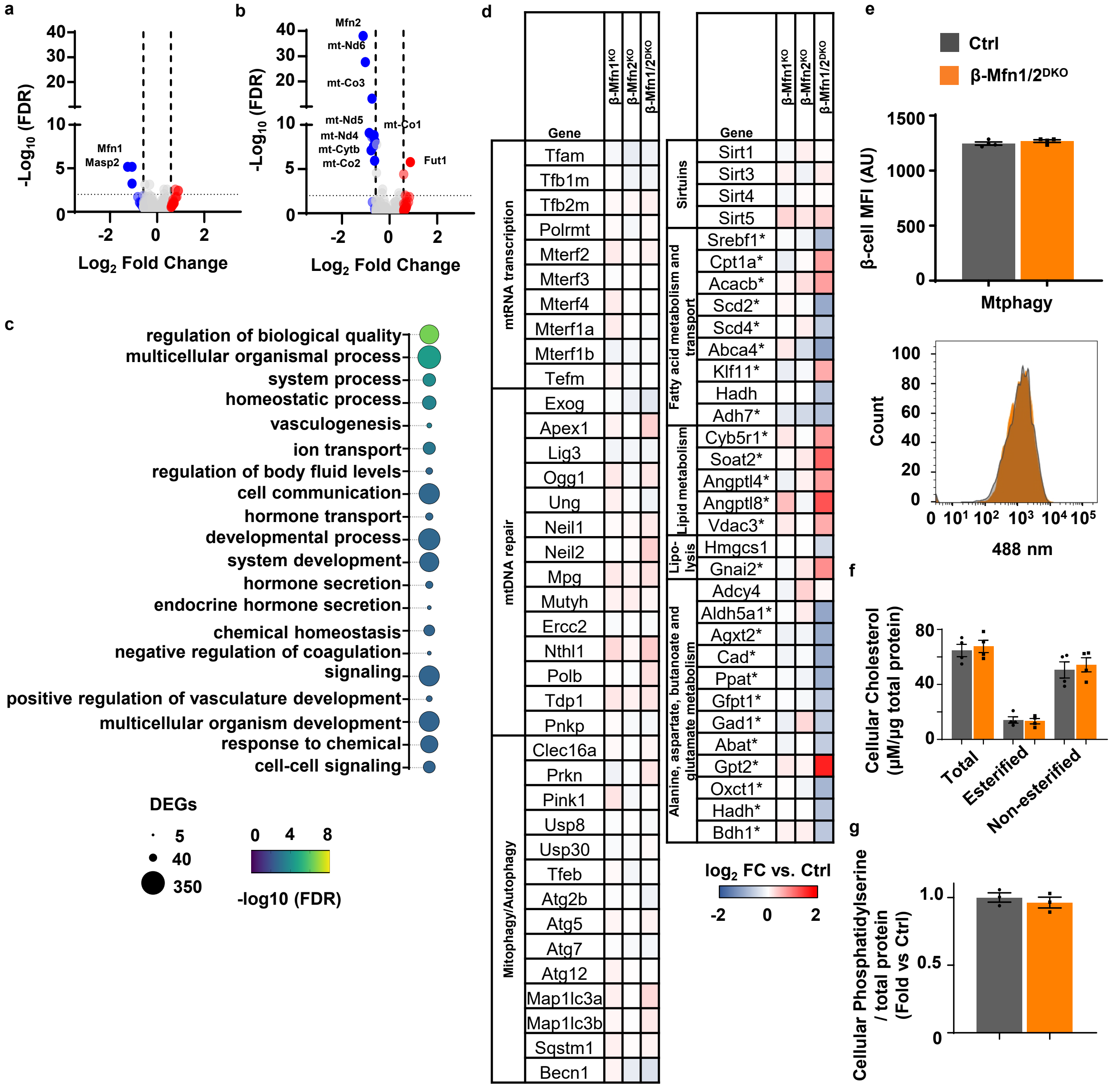


**Figure S4. Transcriptomic profiling suggests defects in specific metabolic pathways, while mtDNA repair, mitophagy, total/esterified cholesterol, and phosphatidylserine concentrations remain unchanged in Mfn1/2-deficient islets.**

**a** Volcano plot depicting differential RNA expression in islets of β-Mfn1$^{KO}$ (n=4) mice compared to littermate Ctrl (n=6) mice. Significantly differentially expressed genes demarcated by -log10 FDR > or < 2 and log2 fold change (FC) > or < 0.6. **b)** Volcano plot depicting differential RNA expression in islets of β-Mfn2$^{KO}$ (n=5) mice compared to littermate Ctrl (n=6) mice. Significantly differentially expressed genes demarcated by -log10 FDR > or < 2 and log2 fold change (FC) > or < 0.6. **c** Gene ontology (GO) enrichment analysis of top differentially regulated GO terms in β-Mfn1/2$^{DKO}$ (n=5) islets compared to littermate Ctrl (n=6) islets, demarcated by both FDR and the number of significantly DEGs. **d** Differential RNA expression heatmap of selected genes from islets of β-Mfn1$^{KO}$ (n=4), β-Mfn2$^{KO}$ (n=5) and β-Mfn1/2$^{DKO}$ (n=5) mice compared to littermate Ctrl mice (n=6). * Benjamini-Hochberg FDR (Padj) <0.05 comparing β-Mfn1/2$^{DKO}$(*) to Ctrl. **e** Assessment of mitophagy by flow cytometric quantification of Mtphagy dye mean fluorescence intensity (MFI - top) and a histogram plot of intensity following excitation at 488 mm (bottom) in β-cells of 6-week-old Ctrl and β-Mfn1/2$^{DKO}$ islets following incubation with 100 nM Mtphagy dye for 3 hours. n=4/group. Data are presented as mean ± SEM. **f** Cellular total cholesterol and cholesteryl ester concentrations (normalized to total protein) in 10-week-old Ctrl and β-Mfn1/2$^{DKO}$ islets. n=4/group. Data are presented as mean ± SEM. **g** Cellular phosphatidylserine levels (normalized to total protein) in 10-week-old Ctrl and β-Mfn1/2$^{DKO}$ islets. n=3/group. Data are presented as mean ± SEM. Source data are provided as a Source Data file.

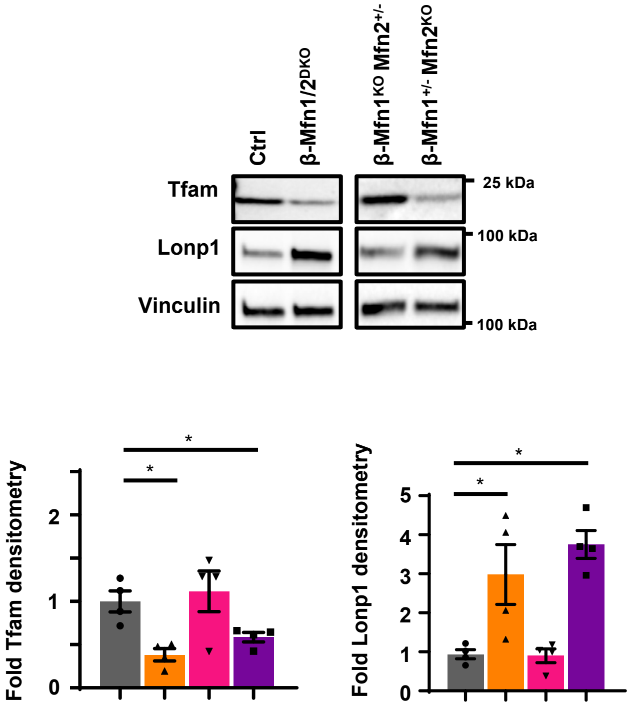


**Figure S5. Tfam and Lonp1 protein expression is preserved in mice bearing a single allele of *Mfn2*, but not *Mfn1*, in β-cells.**

Expression of Tfam and Lonp1 by Western blot (WB) in islets isolated from 11-week-old Ctrl, β-Mfn1/2$^{DKO}$, β-Mfn1$^{KO}$Mfn2$^{+/-}$ and β-Mfn1$^{+/-}$Mfn2$^{KO}$ littermates. Representative of 4 independent mice per group. Tfam (left) and Lonp1 (right) densitometry, normalized to vinculin. n=4/group; Data are presented as mean ± SEM. *p=0.026 for for Ctrl vs β-Mfn1/2$^{DKO}$, p=0.0219 for Ctrl vs β-Mfn1$^{+/-}$Mfn2$^{KO}$ Tfam densitometry, p=0.0184 for Ctrl vs β-Mfn1/2$^{DKO}$, p=0.002 for Ctrl vs β-Mfn1$^{+/-}$Mfn2$^{KO}$ Lonp1 densitometry. All statistical analyses were performed by one-way ANOVA followed by Sidak's post-hoc test for multiple comparisons. Of note, studies in Ctrl and β-Mfn1/2$^{DKO}$ mice were performed together alongside β-Mfn1$^{KO}$, β-Mfn2$^{KO}$, β-Mfn1$^{KO}$Mfn2$^{+/-}$ and β-Mfn1$^{+/-}$Mfn2$^{KO}$ littermates, and thus densitometry appears twice for purposes of relevant comparisons (in Figure 6B and again in Figure S5). Uncropped western blots and source data are provided as a Source Data file.

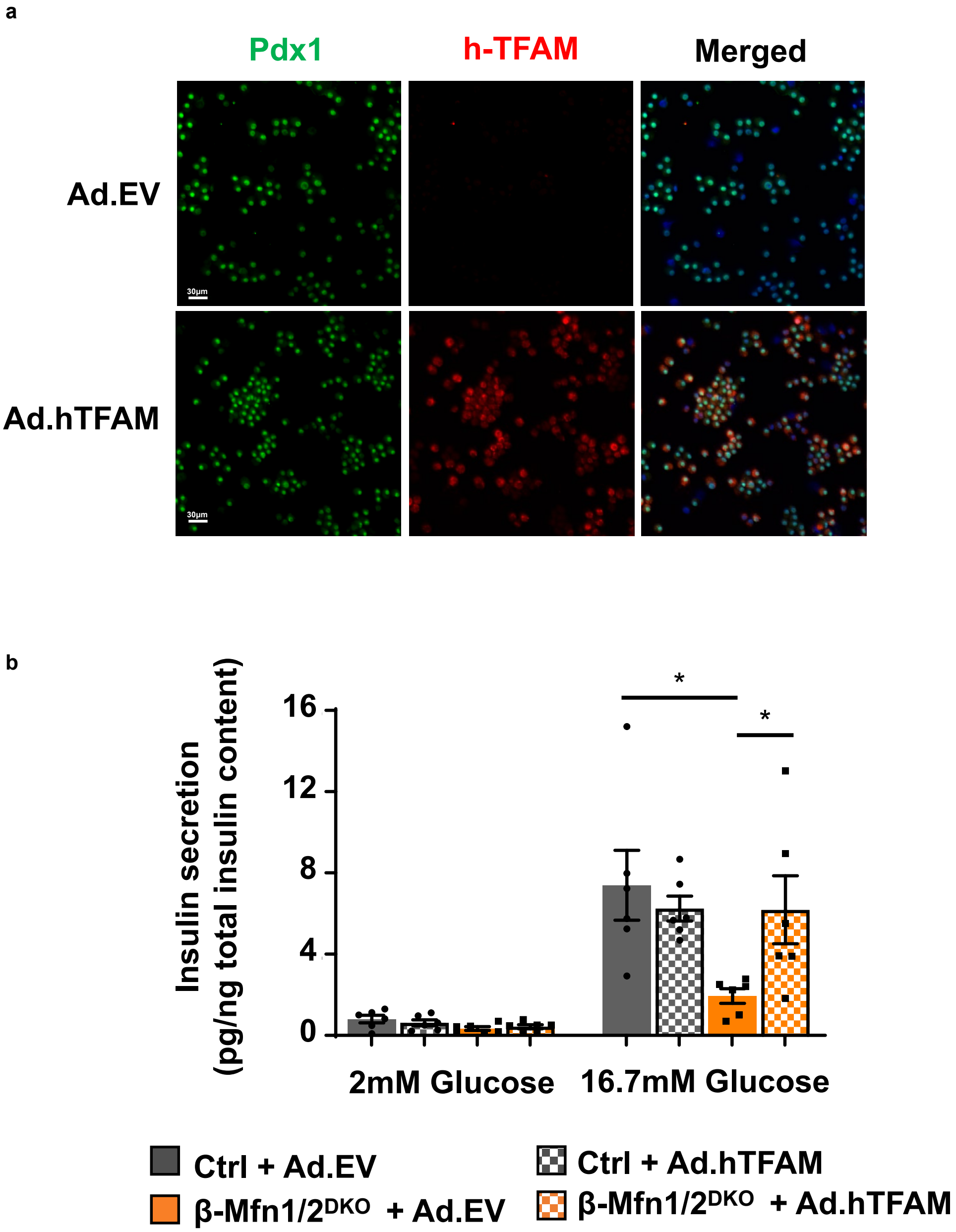


**Figure S6. Overexpression of human-specific TFAM in primary mouse islets.**
**a** Immunofluorescence image at 20X magnification of mouse islets following transduction with Ad.EV or Ad.hTFAM adenoviral particles. Prior to fixation and staining, transduced islets were dispersed into single cells and cytocentrifugated onto glass slides. Samples stained for Pdx1 (green), human-specific TFAM (red), and DAPI (blue). Representative of 3 independent experiments. **b** Glucose-stimulated insulin secretion (normalized to total islet insulin content) following static incubations in 2mM and 16.7 mM glucose, performed in isolated Ctrl and β-Mfn1/$2^{DKO}$ islets following transduction with Ad.EV or Ad.hTFAM adenoviral particles. n=6/group. Data are presented as mean ± SEM. *p=0.0005 for Ctrl + Ad.EV vs β-Mfn1/2DKO + Ad.EV, p=0.0083 for β-Mfn1/2DKO + Ad.EV vs β-Mfn1/2DKO + Ad.hTFAM by one-way ANOVA followed by Tukey's post-hoc test for multiple comparisons. Source data are provided as a Source Data file.

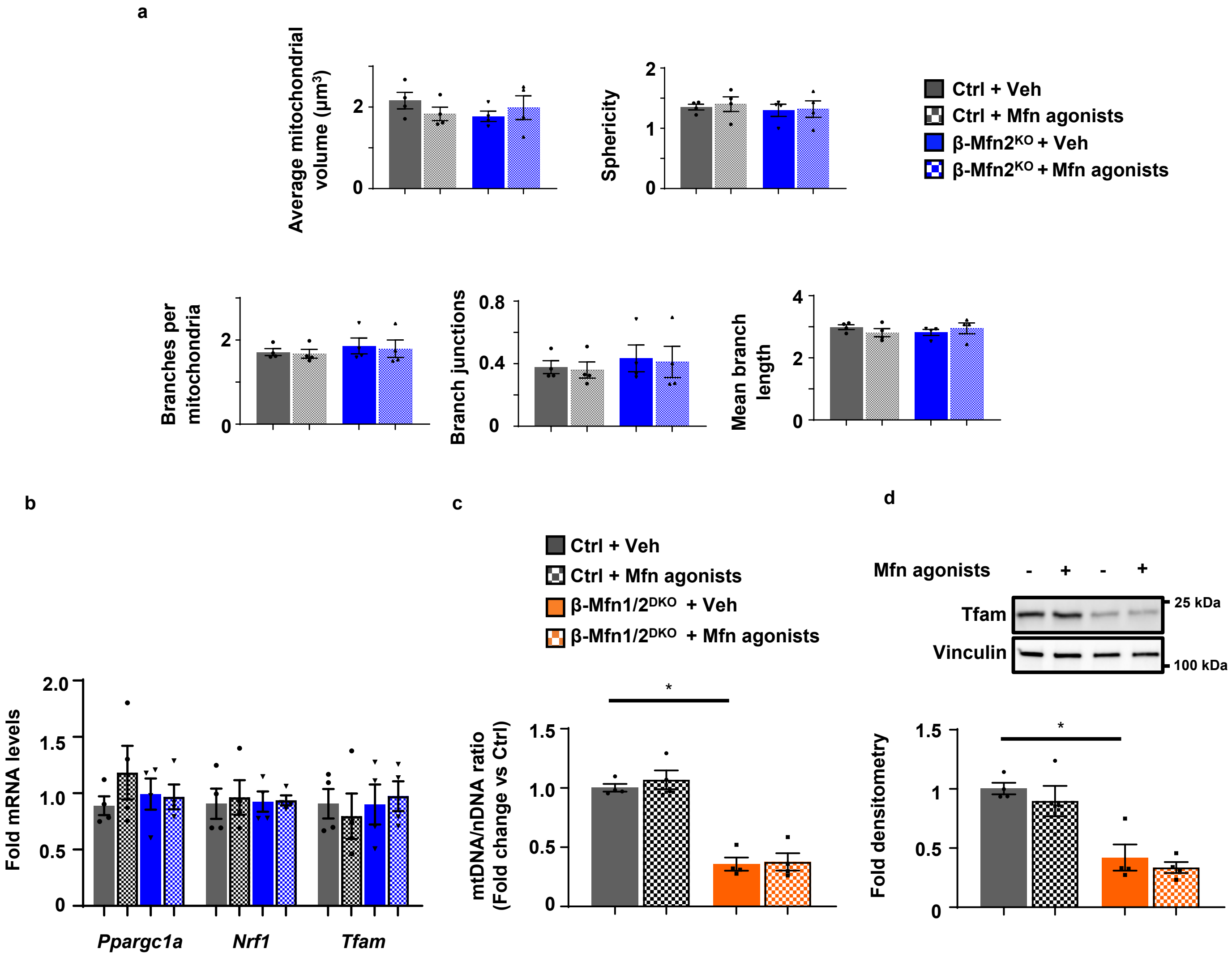


**Figure S7. Effects of Mfn agonists on mitochondrial morphology, mitochondrial biogenesis, mtDNA content, and Tfam levels in mitofusin-deficient islets.**

**a** β-cell mitochondrial morphology and network analysis by MitoAnalyzer of deconvolution immunofluorescence Z-stack images, stained for SDHA (and insulin), from pancreatic islets of Ctrl and β-Mfn2$^{KO}$ mice following treatment with vehicle (Veh) or 0.5 μM Mfn agonists for 24 hrs. n=4/group. Data are presented as mean ± SEM. **b** Relative mRNA expression of *Ppargc1a*, *Nrf1*, and *Tfam* by RT-qPCR (normalized to *HPRT* expression) from isolated islets of Ctrl and β-Mfn1/2$^{DKO}$ mice following treatment with vehicle (Veh) or 0.5 μM Mfn agonists for 24 hrs. n=4/group. Data are presented as mean ± SEM. **c** Relative mtDNA content measured by qPCR (normalized to nuclear DNA expression) from isolated islets of Ctrl and β-Mfn1/2$^{DKO}$ mice following treatment with vehicle (Veh) or 0.5 μM Mfn agonists for 24 hrs. n=4/group. Data are presented as mean ± SEM. *p<0.0005 by one-way ANOVA followed by Tukey's post-hoc test for multiple comparisons. **d** Expression of Tfam by Western blot (WB; top) from isolated islets of Ctrl and β-Mfn1/2$^{DKO}$ mice following treatment with vehicle (Veh) or 0.5 μM Mfn agonists for 24 hrs. Tfam (bottom) densitometry, normalized to vinculin. n=4/group. Data are presented as mean ± SEM. *p=0.0028 by ANOVA followed by Sidak's post-hoc test for multiple comparisons. Uncropped westerns and source data are provided as a Source Data file.

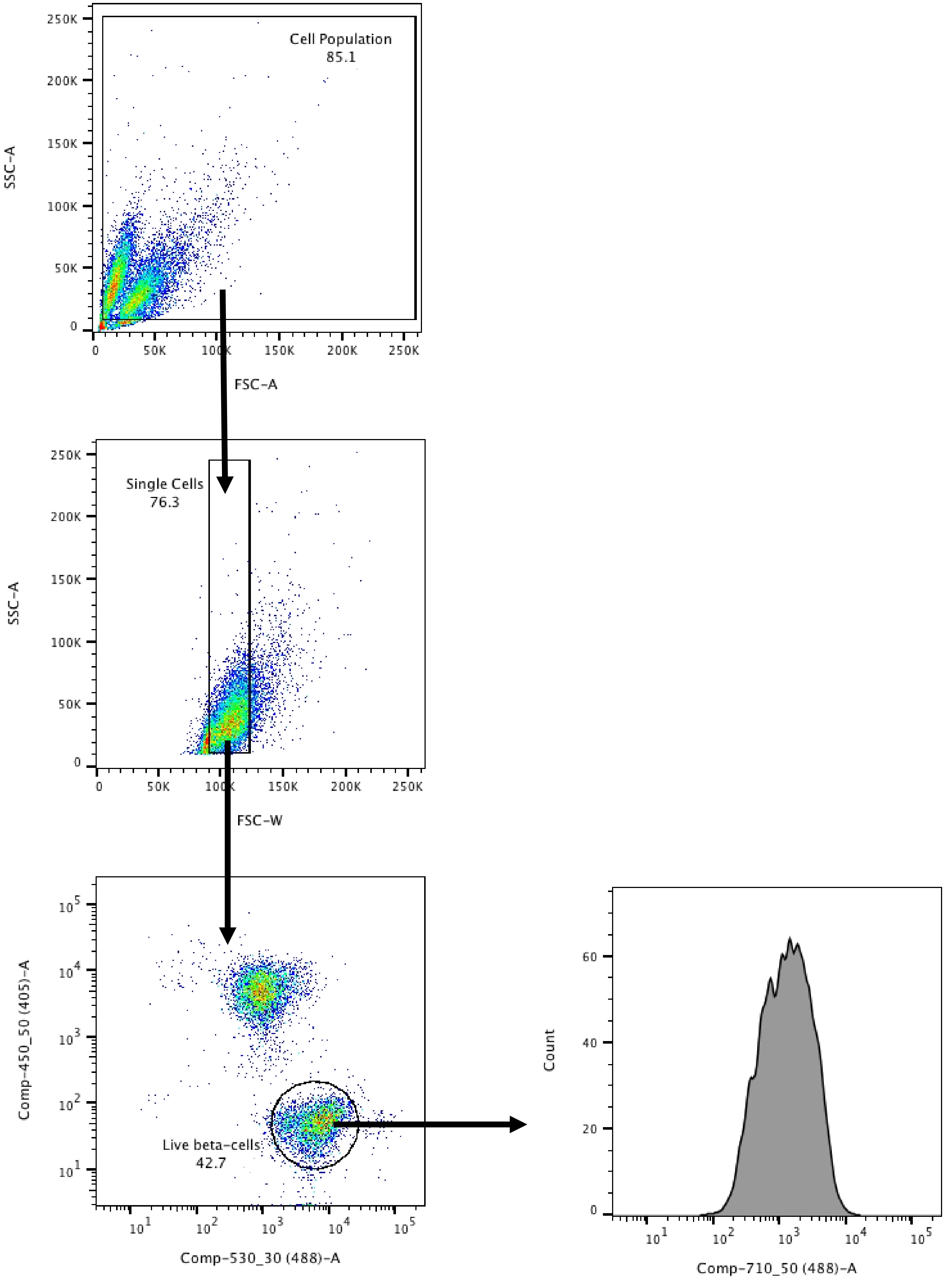


**Figure S8. Flow Cytometry gating strategy:** Representative plots generated from dissociated mouse islets analyzed by flow cytometry. Cells were first gated using FSC-A and SSC-A plots to exclude cell debris. Single cells were then gated using FSC-W and SSC-A plots to exclude cell clumps. DAPI and Fluozin-3 staining was then used to analyze live β-cell populations. Median fluorescence intensity for MtPhagy dye was then calculated for Ctrl and β-Mfn1/2$^{DKO}$ islets as represented in Figure S4E.

Figure S1D

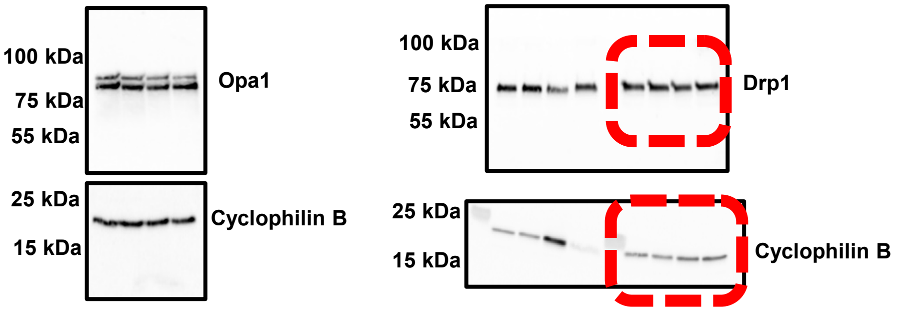


Figure S5

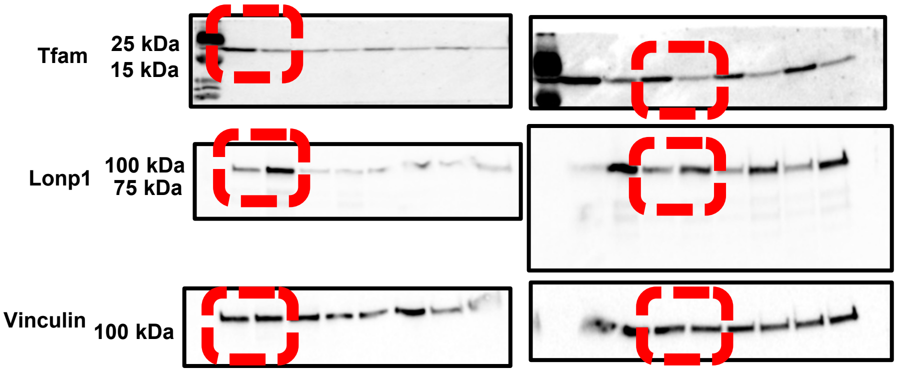


Figure S7D

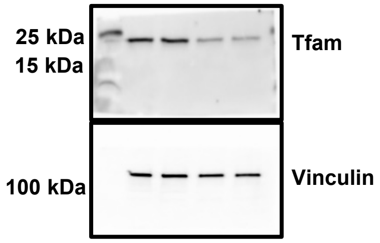


**Figure S9. Uncropped western blots for Supplementary figures S1D, S5, S7D.**
